# Predictive modeling reveals that higher-order cooperativity drives transcriptional repression in a synthetic developmental enhancer

**DOI:** 10.1101/2021.07.28.454075

**Authors:** Yang Joon Kim, Kaitlin Rhee, Jonathan Liu, Paul Jeammet, Meghan Turner, Stephen Small, Hernan G. Garcia

## Abstract

A challenge in quantitative biology is to predict output patterns of gene expression from knowledge of input transcription factor patterns and from the arrangement of binding sites for these transcription factors on regulatory DNA. We tested whether widespread thermodynamic models could be used to infer parameters describing simple regulatory architectures that inform parameter-free predictions of more complex enhancers in the context of transcriptional repression by Runt in the early fruit 2y embryo. By modulating the number and placement of Runt binding sites within an enhancer, and quantifying the resulting transcriptional activity using live imaging, we discovered that thermodynamic models call for higher-order cooperativity between multiple molecular players. This higher-order cooperativity capture the combinatorial complexity underlying eukaryotic transcriptional regulation and cannot be determined from simpler regulatory architectures, highlighting the challenges in reaching a predictive understanding of transcriptional regulation in eukaryotes and calling for approaches that quantitatively dissect their molecular nature.

## 1 Introduction

During embryonic development, transcription factors bind stretches of regulatory DNA termed enhancers to dictate the spatiotemporal dynamics of gene expression patterns that will lay out the future body plan of multicellular organisms [Spitz and Furlong, 2012, Small and Arnosti, 2020]. One of the greatest challenges in quantitative developmental biology is to predict these patterns from knowledge of the number, placement, and affinity of transcription factor binding sites within enhancers. The early embryo of the fruit fly *Drosophila melanogaster* has become one of the main workhorses in this attempt to achieve a predictive understanding of cellular decision-making in development due to its well-characterized gene regulatory network and transcription factor binding motifs, and the ease with which its development can be quantified using live imaging [Garcia et al., 2020, Small and Arnosti, 2020, Rivera et al., 2019].

Predictive understanding calls for the derivation of theoretical models that generate quantitative and experimentally testable predictions. Thermodynamic models based on equilibrium statistical mechanics have emerged as a widespread theoretical framework to achieve this goal [Ackers et al., 1982, Vilar and Leibler, 2003, Bolouri and Davidson, 2003, Bintu et al., 2005b, a, Segal et al., 2008, Fakhouri et al., 2010, Sayal et al., 2016, Phillips et al., 2019, Eck et al., 2020]. For instance, over the last decade, a dialogue between these thermodynamic models and experiments demonstrated the capacity to quantitatively predict bacterial transcriptional regulation from knowledge of the DNA regulatory architecture [He et al., 2010, Garcia and Phillips, 2011, Brewster et al., 2014, Garcia et al., 2012, Sepulveda et al., 2016].

The predictive power of these models is evident when inferring model parameters from simple regulatory architectures [Boedicker et al., 2013a,b, Razo-Mejia et al., 2018, Phillips et al., 2019]. Consider, for example, that RNA polymerase II (RNAP)—which we take as a proxy for the whole basal transcriptional machinery—binds to a promoter with a dissociation constant *K_p_*. When RNAP is bound, transcription is initiated at a rate *R* (Fig. 1A). In the absence of any regulation, a thermodynamic model will only have *K_p_* and *R* as its free parameters which can be experimentally determined by, for example, measuring mRNA distributions [Razo-Mejia et al., 2020]. Now, we assume that the parameters *K_p_* and *R* inferred in this step do not just enable a 1t to the data, but that their values represent physical quantities that remain unaltered as more complex regulatory architectures are iteratively considered. As a result, when we consider the case where a single repressor molecule can bind, our model calls for only two new free parameters: a dissociation constant for repressor to its binding motif *K_r_*, and a negative cooperativity between repressor and RNAP, *ω_rp_*, that makes the recruitment of RNAP less favorable when the repressor is bound to its binding site (Fig. 1B). Once again, after determining *K_r_* and *ω_rp_* experimentally [Phillips et al., 2019], we consider the case where two repressors can bind simultaneously (Fig. 1C). If the repressors interact with RNAP independently of each other, then our model has no remaining free parameters such that we will have reached complete predictive power. However, protein-protein interactions between repressors could exist or even higher-order interactions giving rise to a repressor-repressor-RNAP ternary complex might be present. The extra complexity represented by these interactions would require yet another round of experimentation to quantify these interactions represented by *ω_rr_* and *ω_rrp_* in Figure 1C, respectively. Even after quantifying these parameters, predictive power might not be reached if, after adding yet another repressor binding site, a complex between all three repressors and RNAP can be formed (Fig. 1D).

**Figure 1.**
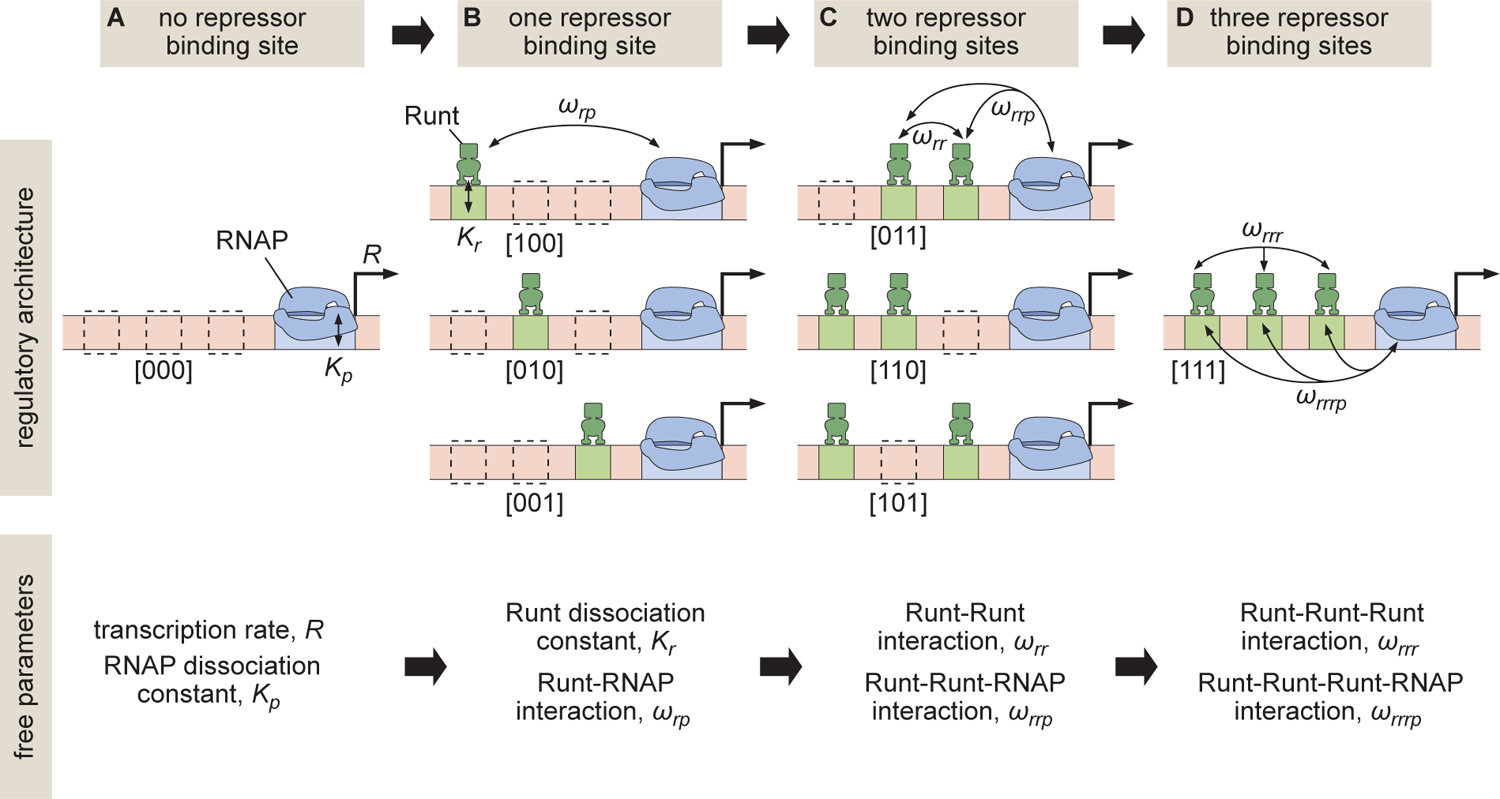
Building up predictive models of transcriptional repression. **(A)** In the absence of repressor binding, gene expression can be characterized by a dissociation constant between RNAP and the promoter *K_p_* and the rate of transcription initiation when the promoter is bound by RNAP *R*. **(B)** In the presence of a single repressor binding site, models need to account for two additional parameters describing the repressor dissociation constant *K_r_* and a repressor-RNAP interaction term *ω_rp_*. **(C)** For two-repressor architectures, parameters accounting for repressor-repressor interactions *ω_rr_* and for interactions giving rise to a repressor-repressor-RNAP complex could also have to be incorporated. **(D)** For the case of three repressor binding sites, additional parameters *ω_rrr_* and *ω_rrrp_* capturing the higher-order cooperativity between three repressor molecules and between three Runt molecules and RNAP, respectively, could be necessary. Note the nomenclature shown below each construct, which indicates which Runt binding sites are present in each construct.

While protein-protein cooperativity captured by *ω_rr_* has been studied both in bacteria [*Ackers et al., 1982*, *Ptashne and Gann, 2002*] and eukaryotes [*Giniger and Ptashne, 1988*, *Ma et al., 1996*, *Lebrecht et al., 2005*, *Parker et al., 2011*, *Fakhouri et al., 2010*, *Sayal et al., 2016*], the necessity of accounting for the higher-order interactions such as those described in our example by the *ω_rrp_* and *ω_rrrp_* terms had only been demonstrated in archeae [*Peeters et al., 2013*] and bacteria [*Dodd et al., 2004*]. The need to invoke this higher-order cooperativity in eukaryotes only became apparent in the last few years [*Estrada et al., 2016b*, *Park et al., 2019*, *Biddle et al., 2020*]. These higher-order cooperativities might be necessary in order to account for the complex interactions mediated by, for example, the recruitment of co-repressors [*Courey and Jia, 2001*, *Walrad et al., 2011*], mediator complex [*Park et al., 2019*], or any other element of the transcriptional machinery. As a result, while posing a challenge to reaching a parameter-free predictive understanding of transcriptional regulation, higher-order cooperativity provides an avenue for quantifying the complexity of the molecular processes underlying eukaryotic cellular decision-making.

In this paper, we sought to test whether an iterative and predictive approach, such as that outlined in Figure 1, was possible for transcriptional repression in the early embryo of the fruit fly *Drosophila melanogaster* or whether it is necessary to invoke higher-order cooperativities that challenge the reach of our predictive models as we add more complexity to the system. To make this possible, we engineered binding sites for the Runt repressor into the Bicoid-activated *hunchback* P2 minimal enhancer. We systematically varied the number and placement of Runt binding sites within this enhancer [Chen et al., 2012] in order to determine whether model 1ts to real-time transcriptional measurements from the enhancer constructs containing only one-Runt binding site could accurately predict repression in two- and three-Runt binding site constructs (Fig. 1A and B). We found that a thermodynamic model can recapitulate all our data. However, we also discovered that, while the model could describe repression by a single Runt repressor, protein-protein and higher-order cooperativities had to be invoked in order to quantitatively account for regulation by two or more repressor molecules. While these higher-order cooperativities limit the iterative bottom-up discourse between theory and experiment that has been successful in bacteria [Phillips et al., 2009], they also provide a concrete theoretical framework for quantifying the complexities behind eukaryotic transcriptional control, and calling for the development of new theories and experiments specifically conceived to uncover the the molecular underpinnings of this complexity.

## 2 Results

### 2.1 Predicting transcription rate using a thermodynamic model of Bicoid activation and Runt repression

We built a predictive model of Runt repression on the Bicoid-activated *hunchback* P2 enhancer using the thermodynamic model framework [Phillips et al., 2019, Bintu et al., 2005b,a] with the goal of predicting the rate of transcription initiation as a function of input transcription factor concentration, and the number and placement of Runt repressor binding sites. Our model rests on the “occupancy hypothesis” that states that the rate of mRNA production, *d*[*mRNA*]/*dt*, is proportional to the probability of the promoter being bound by RNA polymerase II (RNAP), *p_bound_*, such that

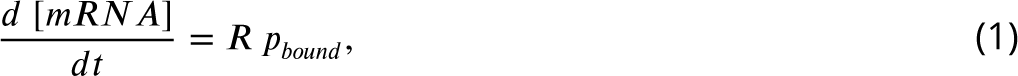

where *R* is the rate of mRNA production when the promoter is occupied by RNAP. Note that, throughout this study, we treat the rate of transcription initiation and the rate of RNAP loading interchangeably.

To generate intuition, we start by modeling the case of *hunchback* P2 with one Runt binding site. Figure 2A illustrates the possible states the system can be found in. Each state has an associated statistical weight which can be calculated as prescribed by equilibrium statistical mechanics [Bintu et al., 2005b,a]. Here, we assume that there are six Bicoid binding sites with the same dissociation constant given by *K_b_*, one Runt binding site with a dissociation constant specified by *K_r_*, and a promoter with a dissociation constant for RNAP prescribed by *K_p_*. In the absence of Runt, we consider four states as shown in the top two rows of Figure 2A. Here, we assume that Bicoid-Bicoid cooperativity is so strong that the enhancer can either be unoccupied or completely bound by Bicoid molecules [Gregor et al., 2007, Park et al., 2019]. Further, we consider an interaction between Bicoid and RNAP given by *ω_bp_*. For simplicity, we use the dimensionless parameters *b* = [*Bicoid*]/*K_b_*, *r* = [*Runt*]/*K_r_* and *p* = [*RNAP*]/*K_p_*. These assumptions lead to a functional form reminiscent of a Hill function that explains the sharp step-like expression pattern along the embryo’s anterior-posterior axis of the *hunchback* gene [Gregor et al., 2007, Park et al., 2019, Driever and Nusslein-Volhard, 1988, 1989]. A full thermodynamic model in which we do not make this assumption of high Bicoid-Bicoid cooperativity is discussed in detail in Section S1 and Section S2.

**Figure 2.**
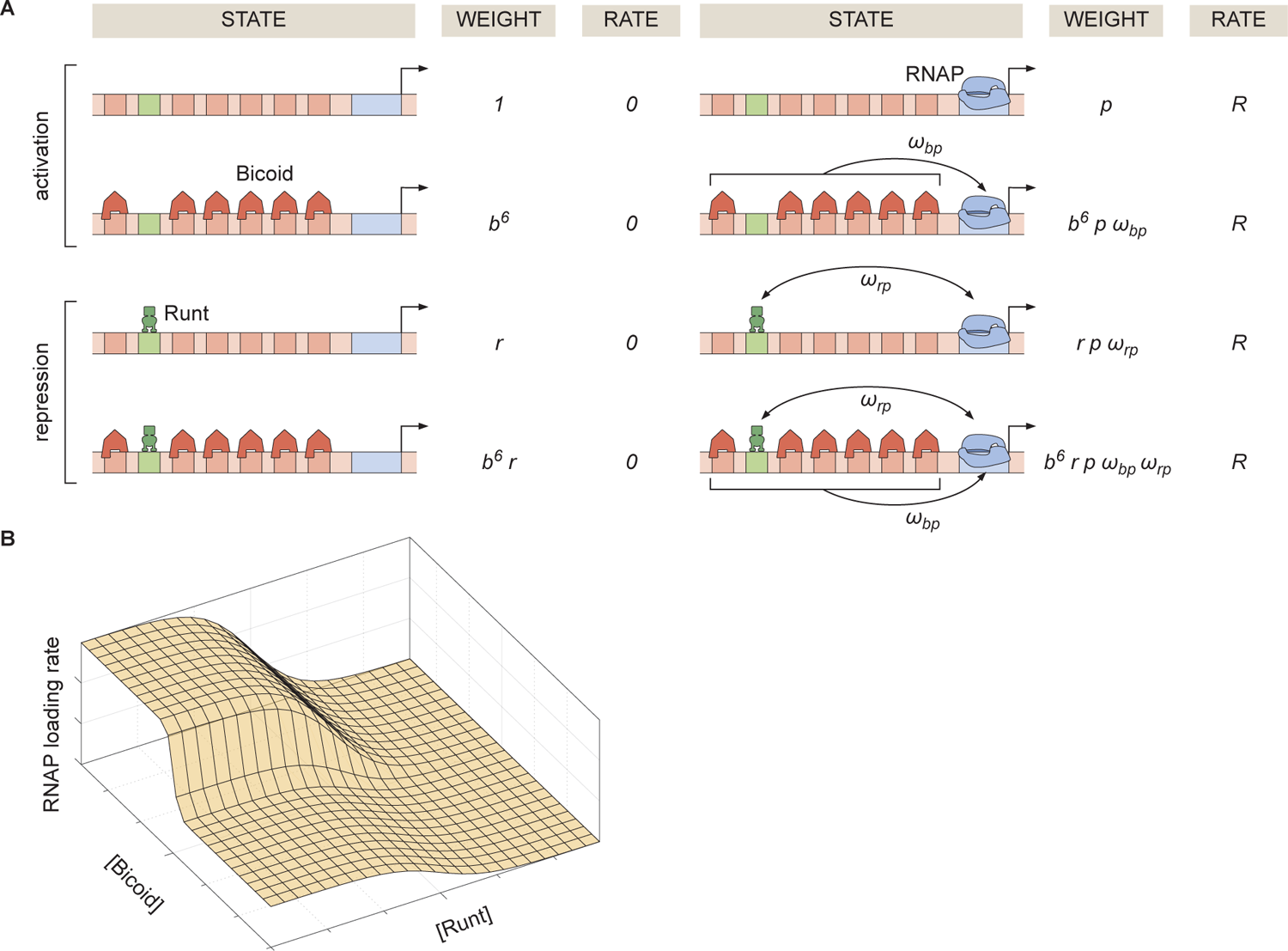
Thermodynamic model of transcriptional regulation by Bicoid activator and Runt repressor. **(A)** States and statistical weights for the regulation of *hunchback* P2 with one Runt binding site in the limit of strong Bicoid-Bicoid cooperativity. Here, we use the dimensionless parameters *b* = [*Bicoid*]/*K_b_*, *r* = [*Runt*]/*K_r_*, and *p* = [*RNAP*]/*K_p_*, where *K_b_*, *K_r_*, and *K_p_* are the dissociation constants of Bicoid, Runt, and RNAP, respectively. *ω_bp_* represents the cooperativity between Bicoid and RNAP, *ω_rp_* captures the cooperativity between Runt and RNAP, and *R* represents the rate of transcription when the promoter is occupied by RNAP. The top two rows correspond to states where only Bicoid and RNAP act, while the bottom two rows represent repression by Runt. **(B)** Representative prediction of RNAP loading rate as a function of Bicoid and Runt concentrations for *ω_bp_* = 3*, ω_rp_* = 0.001*, p* = 0.001*, R* = 1(*AU* /*min*).

The molecular mechanism by which Runt downregulates transcription of its target genes remains unclear [Chen et al., 2012, Hang and Gergen, 2017, Koromila and Stathopoulos, 2017, 2019]. Here, we assume the so-called “direct repression” model [Gray et al., 1994] that posits that Runt operates by inhibiting RNAP binding to the promoter through a direct Runt-RNAP interaction term given by *ω_rp_ <* 1 independently of Bicoid. As a result, in the presence of Runt, we consider four additional states as shown in the bottom two rows of Figure 2A. Other potential mechanisms of Runt repression are further discussed in Supplementary Section S5), where we also show that the choice of specific mechanism does not change our conclusions.

Given these assumptions, we arrive at the microstates and corresponding statistical weights shown in Figure 2A. The probability of finding RNAP bound to the promoter, *p_bound_*, is calculated by dividing the sum of all statistical weights featuring RNAP by the sum of the weights of all possible microstates.

The calculation of *p_bound_* combined with Equation 1 leads to the expression

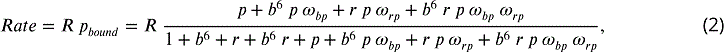

which makes it possible to predict the output rate of mRNA production as a function of the input concentrations of Bicoid and Runt (Fig. 2B). With this theoretical framework in hand, we experimentally tested the predictions of this model.

### 2.2 Measuring transcriptional input-output to test model predictions

The transcriptional input-output function in Figure 2B indicates that, in order to predict the rate of RNAP loading and to test our theoretical model, we need to first measure the concentration of the input Bicoid and Runt transcription factors. In order to quantify the concentration profile of Bicoid, we used an established eGFP-Bicoid line [Gregor et al., 2007] and measured mean Bicoid nuclear concentration dynamics along the anterior-posterior axis of the embryo over nuclear cycles 13 and 14 (nc13 and nc14, respectively) as shown in Movie S1 [Eck et al., 2020]. An example snapshot and time trace of Bicoid nuclear concentration dynamics at 40% of the embryo length appear in Figure 3A and B.

**Figure 3.**
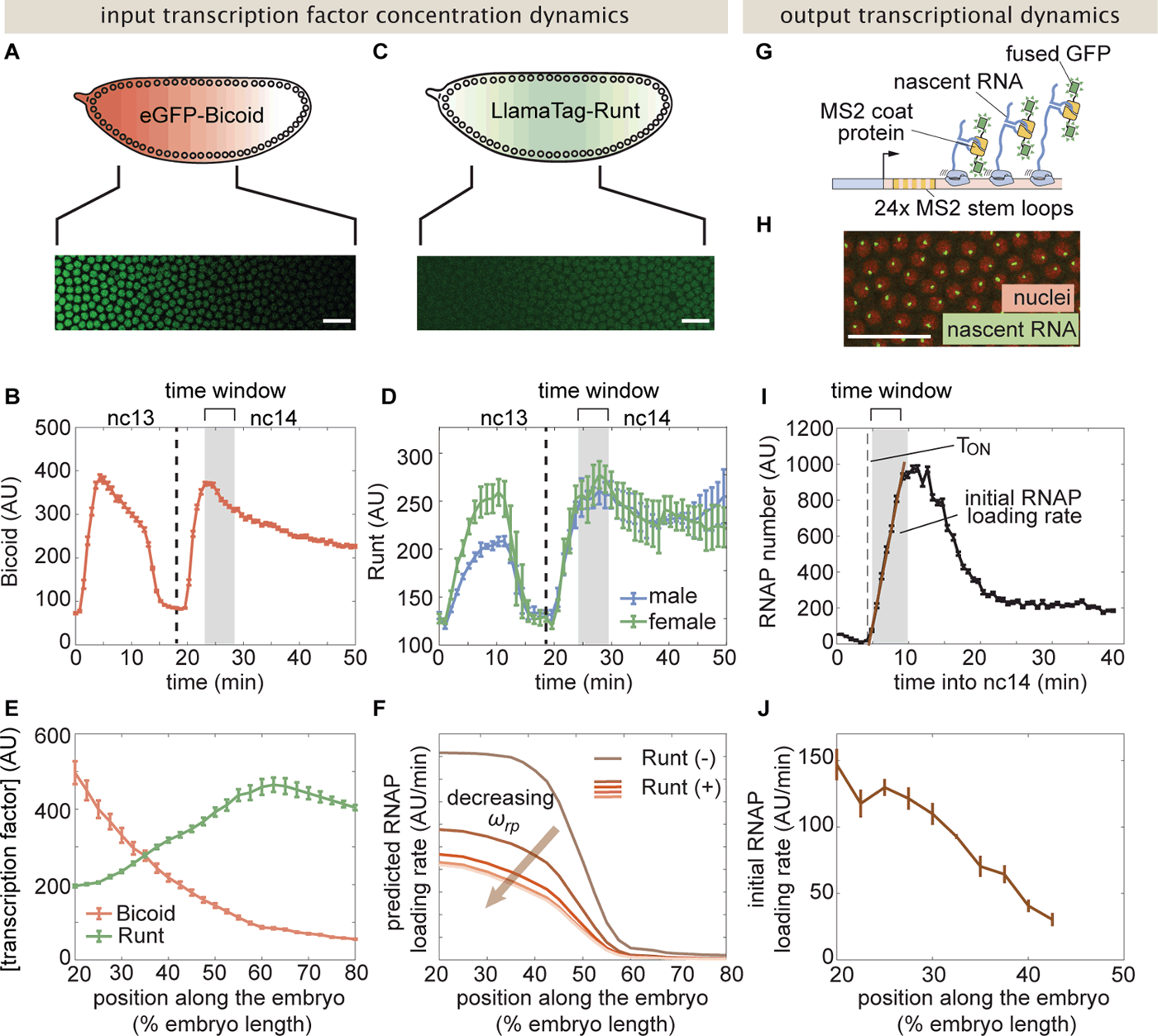
Measurement of input transcription factor concentrations and output rate of transcription to test model predictions. **(A)** Snapshot of an embryo expressing eGFP-Bicoid spanning 20-60% of the embryo length. (For a full time-lapse movie, see Movie S1.) **(B)** Bicoid nuclear fluorescence dynamics taken at 40% of the embryo. **(C)** Snapshot of an embryo expressing eGFP:LlamaTag-Runt spanning 20-60% of the embryo length. (For a full time-lapse movie, see Movie S2.) **(D)** Runt nuclear concentration dynamics in males and females. **(E)** Measured transcription factor concentration profiles along the anterior-posterior axis of the embryo. The concentration profiles are averaged over the gray shaded regions shown in (B) and (D) which corresponds to a time window between 5 and 10 minutes into nc14. **(F)** Predicted RNAP loading rate for *hunchback* P2 with one Runt binding site over the anterior-posterior axis generated for a reasonable set of model parameters *K_b_* = 30 AU, *K_r_* = 100 AU, *ω_bp_* = 100, *p* = 0.001, and *R* = 1 AU/min for varying values of the Runt-RNAP interaction term *ω_rp_* = [10^−2^, 1]. **(G)** Schematic of the MS2 system where 24 repeats of the MS2 loop sequence are inserted downstream of the promoter followed by the *lacZ* gene. The MS2 coat protein (MCP) fused to GFP binds the MS2 loops. **(H)** Example snapshot of an embryo expressing MCP-GFP and Histone-RFP. Green spots to active transcriptional loci and red circles correspond to nuclei. Spot intensities are proportional to the number of actively transcribing RNAP molecules. **(I)** Representative MS2 fluorescence averaged over a narrow window (2.5% of the embryo length) along the anterior-posterior axis of the embryo. The initial rate of RNAP loading was obtained by 1tting a line (brown) to the initial rise of the data. **( J)** Measured initial rate of RNAP loading (over a spatial bin of 2.5% of the embryo length) across the anterior-posterior axis of the embryo, from the *hunchback* P2 enhancer. (B, D, E, and J, error bars represent standard error of the mean over 3 embryos; I, error bars represent standard error of the mean over the spatial averaging corresponding to roughly ten nuclei; A, C, and H, white scale bars represent 20 *µ*m.)

Quanti1cation of the Runt concentration using standard 2uorescent protein fusions is not possible due to the slow maturation times of these proteins [Bothma et al., 2018]. We therefore measured Runt concentration dynamics using our recently developed LlamaTags, which are devoid of such maturation dynamics artifacts [Bothma et al., 2018]. Speci1cally, we generated a new fly line harboring a fusion of a LlamaTag against eGFP to the endogenous *runt* gene using CRISPR/Cas9-mediated homology-directed repair (Materials and Methods; Harrison et al. [2010], Gratz et al. [2015]).

Using this LlamaTag fusion, we measured the mean Runt nuclear fluorescence along the anterior-posterior axis of the embryo over nc13 and nc14 (Materials and Methods; Figure 3B; Movie S2). As expected due to the location of the *runt* gene on the X chromosome [Lott et al., 2011], there is a sex dependence in the nuclear concentration levels in nc13, with males displaying lower Runt levels than females; this difference is compensated by early nc14 (Fig. 3C,D). As a result, for ease of analysis, we focused subsequent quantitative dissection on nc14.

We used the measured input protein concentration pro1les to predict the output transcription rate. To make this possible, we invoked previous observations stating that the concentration dynamics of input transcription factors does not signi1cantly affect the initial rate of RNAP loading [Garcia et al., 2013, Eck et al., 2020]. As a result, we decided to use the time-averaged concentration dynamics of Bicoid and Runt over a time window spanning 5 min after the 13th anaphase to 10 min after this anaphase (gray shaded region in Fig. 3B and D) as inputs to our model, resulting in static spatial concentration profiles shown in Figure 3E. We then used these time-averaged concentration profiles of input transcription factors to calculate the time-averaged rate of transcription initiation over the same time window. In the Supplementary Information Section S3 we compare this methodology with one that acknowledges input transcription factor concentration dynamics and show that the prediction stemming from both approaches leads to equivalent theoretical predictions. Notably, the time-averaged rate of transcription predicted by the dynamic inputs was similar to the rate of transcription predicted by the static inputs.

Along the anterior-posterior axis of the embryo, the measured Bicoid and Runt concentration pro1les de1ne a trajectory through the input-output function (Fig. 2B). Given a set of parameters, this trajectory predicts the initial rate of RNAP loading. This quantitative prediction can be directly compared with experimentally measured transcription initiation rates. For example, given the concentration profiles shown in Figure 3E, we calculate the RNAP loading rate as a function of the position along the embryo for different values of the Runt-RNAP interaction, captured by *ω_rp_* (Fig. 3F). As expected, we predict that the rate of transcription decreases as *ω_rp_*, describing Runt-RNAP cooperativity, decreases.

Next, we sought to experimentally test these predictions by measuring the rate of RNAP loading using the MS2 system [Bertrand et al., 1998, Lucas et al., 2013, Garcia et al., 2013]. Here, we inserted 24 repeats of the MS2 loop sequence following the *hunchback* P2 enhancer and *even-skipped* promoter in our reporter construct, which leads to the fluorescent labeling of sites of active transcription in living embryos (Fig. 3G and H; Movie S3). The fluorescence intensity of each MS2 spot is proportional to the number of actively transcribing RNAP molecules [Garcia et al., 2013]. In order to quantify the transcriptional activity reported by MS2, we measured the mean MS2 spot fluorescence over nuclei in a narrow spatial window (Fig. 3I [Garcia et al., 2013, Eck et al., 2020]. To measure the initial rate of RNAP loading, we obtained the slope of the initial rise in the number of actively transcribing RNAP molecules over the same time window used to average input transcription factor concentration (Fig. 3I, brown line). The resulting RNAP loading rate plotted over the anterior-posterior axis is in qualitative agreement with the classic pattern driven by the *hunchback* P2 minimal enhancer (Fig. 3J; Garcia et al. [2013], Chen et al. [2012], Park et al. [2019]).

While we chose the initial rate of transcription as the experimental measurable to confront against our model predictions, the MS2 technique can also report on other dynamical features of transcription such as the time window over which transcription occurs and the fraction of loci that engage in transcription at any point over the nuclear cycle. While these two quantities have been shown to be relevant in shaping gene expression patterns in other regulatory contexts [Garcia et al., 2013, Lammers et al., 2020, Eck et al., 2020, Dufourt et al., 2018, Reimer et al., 2021], we found that the transcription time window was not significantly regulated in the presence of Runt. As described in Section S8, we did 1nd some modulation of the fraction of transcriptionally engaged loci for a subset of our synthetic enhancer constructs but, as we could not detect a clear trend in how this fraction of active loci was modulated, we did not pursue a theoretical dissection of the control of this quantity by Runt.

### 2.3 Enhancer sequence dictates unrepressed transcription rates by determining RNAP-promoter interactions

A major assumption of our theoretical approach is that the model parameters obtained from simple regulatory architectures can be used as inputs for more complex constructs. For instance, we assume that the Runt-independent model parameters for Bicoid and RNAP action—*K_b_*, *ω_bp_*, *p* and *R* (Fig. 2A)—are conserved for all constructs containing Runt binding sites regardless of their number and placement in the enhancer. If model parameters can be shared across constructs, then our model should predict the same profile for the rate of transcription across all synthetic enhancer constructs.

To test this assumption, we measured the initial rate of RNAP loading in all of our reporter constructs, in *runt* null embryos (Materials and Methods). Notably, unrepressed transcription rates varied significantly across synthetic enhancers (Fig. 4A). For example, despite no Runt being present, the [001] construct had almost twice the unrepressed rate of [000].

**Figure 4.**
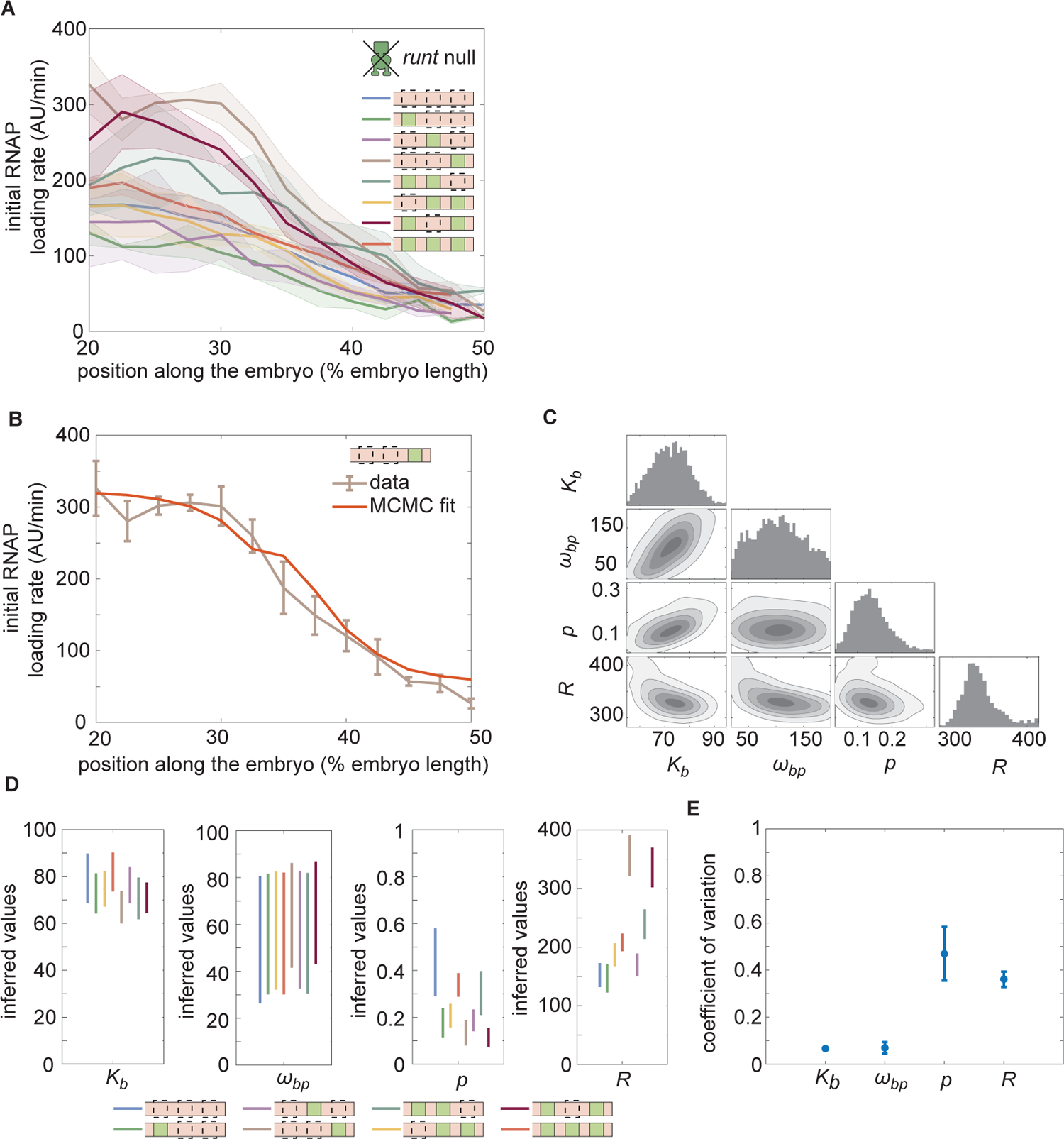
Enchancer-to-enhancer variability in the unrepressed transcription level stems from unique RNAP-dependent parameters. **(A)** Measured initial rates of RNAP loading across the anterior-posterior axis of the embryo for all synthetic enhancer constructs in the *absence* of Runt protein. **(B)** Representative best MCMC 1t and **(C)** associated corner plot for the [001] construct in the *runt* null background. **(D)** Inferred model parameters for all synthetic enhancers in the absence of Runt repressor. **(E)** Coefficient of variation of inferred parameters. (B, C, error bars represent standard error of the mean over >3 embryos; E, error bars represent standard deviations calculated from the MCMC posterior chains; F, error bars are calculated by propagating the standard deviation of individual parameters from their MCMC chains.)

This large construct-to-construct variability in unrepressed transcription rates likely originates from the Runt binding site sequences interfering with some combination of Bicoid and RNAP function. To uncover the mechanistic effect of these Runt binding sites sequences on unrepressed activity, we sought to determine which parameters in our thermodynamic model varied across constructs. In the absence of Runt repressor, only four states remain corresponding to the two top rows of Figure 2A. In this limit, the predicted rate of transcription is given by

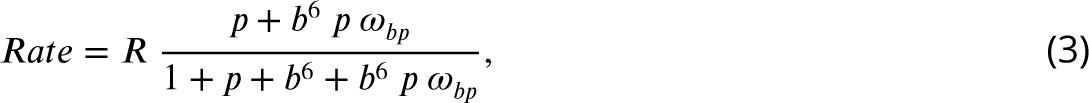

where we have invoked the same parameters as in Equation 2.

To obtain the model parameters for each construct measured in Figure 4A, we used the Bayesian inference technique of Markov Chain Monte Carlo (MCMC) sampling that has been widely used for inferring the biophysical parameters from theoretical models (Liu et al. [2021], Razo-Mejia et al. [2018], Geyer and Thompson [1992]; Supplementary Section S4). A representative comparison of the MCMC 1t to the experimental data reveals good agreement between theory and experiment (Fig. 4B). MCMC sampling also gives the distribution of the posterior probability for each parameter as well as their cross-correlation (Fig. 4C). These corner plots reveal relatively unimodal posterior distributions, suggesting that a unique set of parameters can explain the data.

Note that, while the Bicoid dissociation constant *K_b_* and the Bicoid-RNAP interaction term *ω_bp_* remain largely unchanged regardless of enhancer sequence, there is considerable variability in the inferred mean RNAP-dependent parameters *p* and *R* (Fig. 4D). This variability can be further quantified by examining the coefficient of variation,

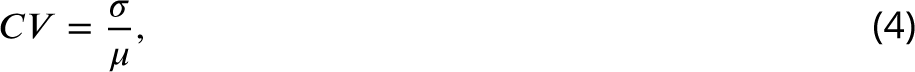

where *σ* and *µ* are the standard deviation and the mean of each parameter, respectively, calculated over all constructs. The coefficients of variation for the RNAP and promoter-dependent parameters are much higher than those for Bicoid-dependent parameters (≈ 40% versus *<* 10%; Fig. 4E). This suggests that the variability in unrepressed transcription rates due to the presence of Runt binding sites is due to differences in the behavior of RNAP at the promoter rather than differences in Bicoid binding or activation being. As a result, as we consider increasingly more complex regulatory architectures, each construct will necessitate its own specific Bicoid- and RNAP-dependent parameters as inferred in Figure 4D. However, we will conserve Runt-dependent parameters as we consider increasingly more complex constructs featuring more Runt binding sites.

### 2.4 The thermodynamic model recapitulates repression by one Runt binding site

Next, we asked whether our model recapitulates gene expression for the *hunchback* P2 enhancer with a one-Runt binding site in the presence of Runt repressor as predicted by Equation 2. We posited that, since the binding site sequence remains unaltered throughout our constructs (Fig. S9), the value of the Runt dissociation constant *K_r_* would also remain unchanged across these enhancers regardless of Runt binding site position; however, we assumed that, as the distance between Runt and the promoter varied, so could the Runt-RNAP interaction term *ω_rp_*.

We measured the initial rate of transcription along the embryo for all our constructs containing one Runt binding site in the presence of Runt protein. We then used MCMC sampling to infer the Runt-dependent parameters *K_r_* and *ω_rp_* for each of these constructs while retaining the mean values of Runt-independent parameters (*K_b_*, *ω_bp_*, *p*, and *R*) obtained from the experiments performed in the absence of Runt (Fig. 4). The resulting MCMC 1ts show signi1cant agreement with the experimental data (Fig. 5A), con1rming that, within our model, the same dissociation constant *K_r_* can be used for all Runt binding sites regardless of their position within the enhancer. Further, the corner plot yielded a unimodal distribution of posterior probability of the inferred parameters (Fig. 5B), indicating the existence of a unique set of most-likely model parameters.

**Figure 5.**
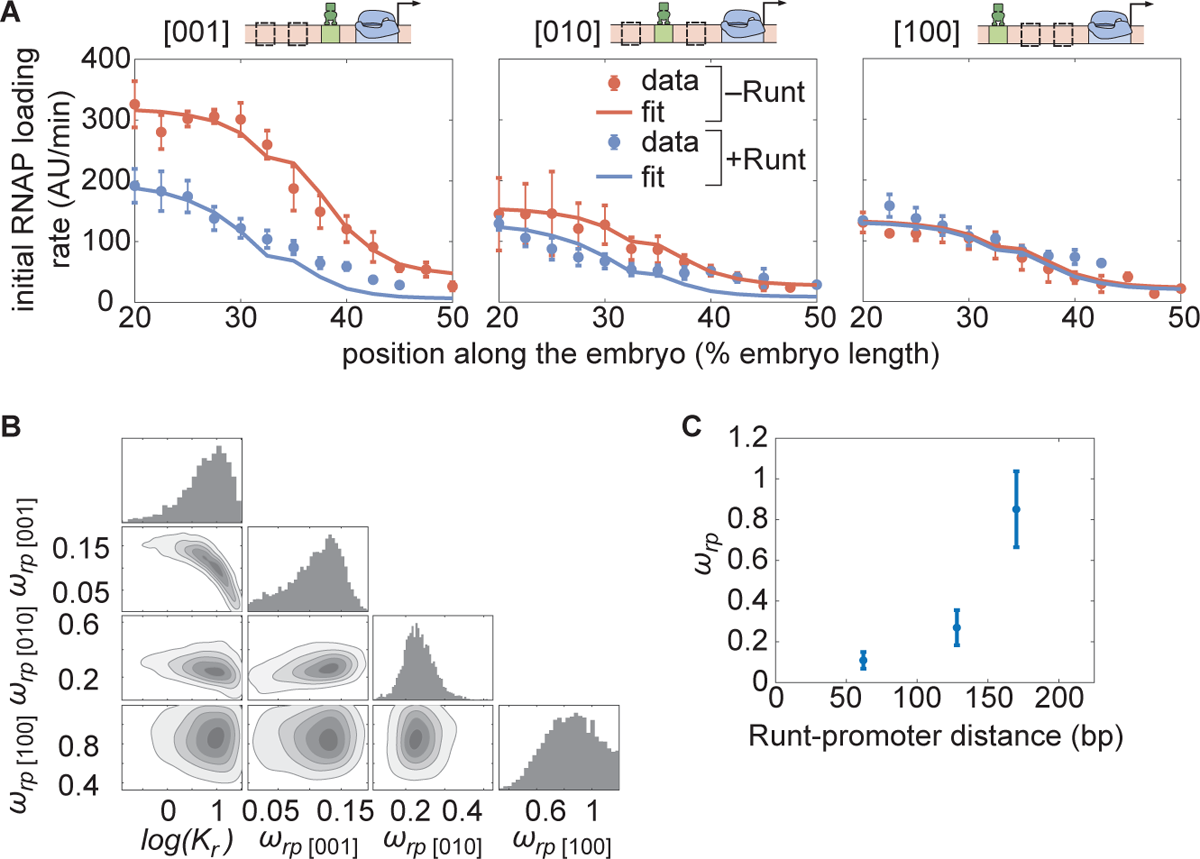
Testing the direct repression model in the presence of one Runt binding site. **(A)** Initial transcription rate as a function of position along the embryo for the three constructs containing one Runt binding site in the presence and absence of Runt repressor, together with their best MCMC 1ts. **(B)** Corner plots from MCMC inference for all constructs with one Runt binding site. **(C)** Inferred *ω_rp_* value as a function of distance between the promoter and the Runt binding site. (B, data points represent mean and standard error of the mean over *>* 3 embryos; D, data and error bars represent the mean and standard deviation of the posterior chains, respectively.)

The observed trend in the Runt-RNAP interaction captured by *ω_rp_* qualitatively agrees with the “direct repression” model. Specifically, because the model assumes that Runt interacts directly with RNAP, it predicts that, the farther apart Runt and the promoter are, the lower this interaction should be [Gray et al., 1994]. In agreement with this prediction, the mean value of *ω_rp_* obtained from our 1ts changes from high repression (*ω_rp_* ≈ 0.1) in the [001] construct to almost no repression (*ω_rp_* ≈ 1) in the [100] construct as the Runt site is moved away from the promoter (Fig. 5C). Thus, the direct repression model recapitulates repression by a single Runt molecule using the the same dissociation constant regardless of Runt binding site position, and displays the expected dependence of the Runt-RNAP interaction term on the distance between these two molecules.

### 2.5 Predicting repression by two-Runt binding sites requires both Runt-Runt and Runt-Runt-RNAP higher-order cooperativity

Could the parameters inferred in the preceding section be used to accurately predict repression in the presence of two Runt binding sites? An extra Runt binding site enables new protein-protein interactions between Runt molecules and RNAP (Fig. 6A). First, we considered individual Runt-RNAP interaction terms, *ω_rp_*_1_ and *ω_rp_*_2_, whose values were already inferred from the one-Runt binding site constructs as *ω_rp_*_[001]_ *, ω_rp_*_[010]_ *, and ω_rp_*_[100]_ (Fig. 5D). Second, we considered protein-protein interactions (positive or negative) between two Runt molecules, *ω_rr_*. Third, following recent studies of Bicoid activation of the *hunchback* P2 minimal enhancer [Estrada et al., 2016a, Park et al., 2019], we also posited the existence of simultaneous Runt-Runt-RNAP higher-order cooperativity *ω_rrp_*. Given these different cooperativities, and as shown in detail in Figure S15B, the predicted rate of transcription is

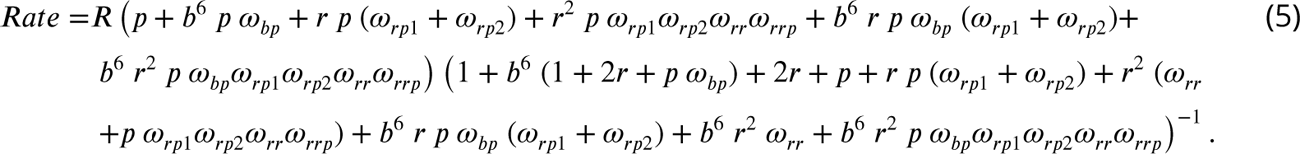

**Figure 6.**
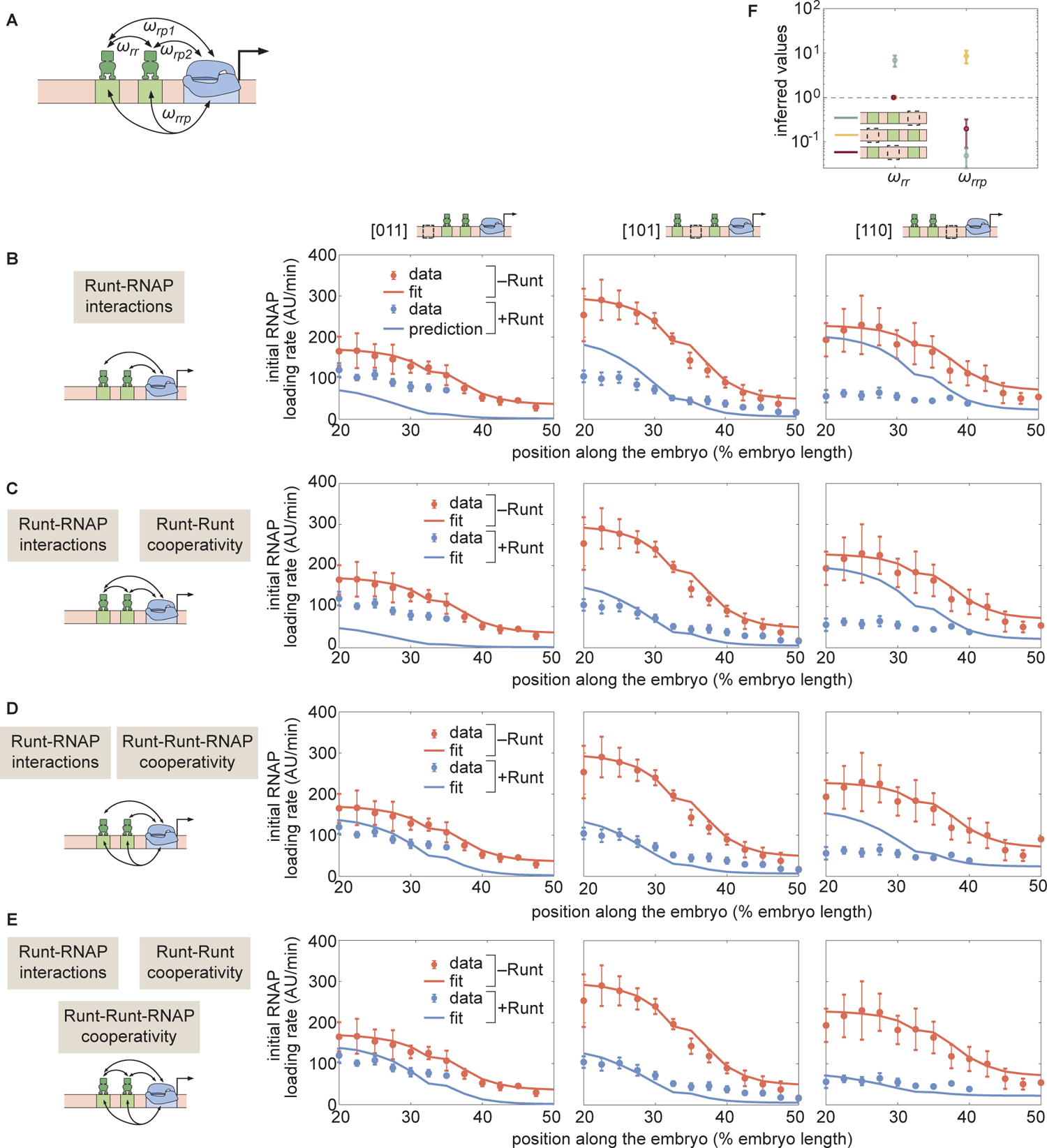
Prediction for the transcription initiation rate of *hunchback* P2 with two-Runt binding sites under different models of cooperativity. **(A)** Direct repression model for *hunchback* P2 with two-Runt binding sites featuring Runt-RNAP interaction terms given by *ω_rp_*_1_ and *ω_rp_*_2_, Runt-Runt cooperativity captured by *ω_rr_*, and Runt-Runt-RNAP higher-order cooperativity accounted for by *ω_rrp_*. **(B)** Parameter-free model prediction for two-Runt binding sites when the two Runt molecules bind the DNA and interact with RNAP independently of each other. **(C,D,E)** Best MCMC 1ts for the data for two-Runt binding site constructs for models with various combinations of cooperativity parameters. **(C)** Model incorporating Runt-Runt cooperativity. **(D)** Model incorporating Runt-Runt-RNAP higher-order cooperativity. **(E)** Model accounting for both Runt-Runt cooperativity and Runt-Runt-RNAP higher-order cooperativity. **(F)** Fixed or inferred parameters *ω_rr_* and *ω_rrp_* for all two-Runt binding site constructs. Note that *ω_rr_* is 1xed to 1 for [011] and [101] constructs due to the fact that no Runt-Runt cooperativity is necessary to quantitatively describe the expression driven by these constructs; only the [110] construct is used to infer both *_rr_* and *ω_rrp_*. The horizontal line of *w* = 1 denotes the case of no cooperativity other than Runt-RNAP cooperativity, *ω_rp_*. (B-E, data points represent mean and standard error of the mean over *>* 3 embryos; F, data and error bars represent the mean and standard deviation of the posterior chain, while the standard deviation for the 1xed *ω_rr_* is set to 0.)

Despite the complexity of this equation, note that its only free parameters are the cooperativity parameters *ω_rr_* and *ω_rrp_*. As a result, we sought to determine whether the Runt-RNAP cooperativity terms, *ω_rp_*_1_ and *ω_rp_*_2_, are sufficient to predict repression by two Runt molecules, or whether the cooperativities given by *ω_rr_* and *ω_rrp_* also need to be invoked.

Consider the simplest case where two Runt molecules bind and interact with RNAP independently from each other. Here, *ω_rr_* = 1, and *ω_rrp_* = 1. This model has no free parameters; all parameters have already been determined by the inferences performed on Runt null datasets and one-Runt binding site constructs (Fig. 4 and Fig. 5, respectively). While there was some agreement between the model and the data for the [101] construct (Fig. 6B, center), significant deviations from the prediction occurred for the other two constructs. These deviations ranged from less repression than predicted for [011] (Fig. 6B, left) to more repression than predicted for [110] (Fig. 6B, right). Thus, this simple model of Runt independent repression is not supported by the experimental data, suggesting additional regulatory interactions between the Runt molecules and RNAP.

A first alternative to the independent repression model is the consideration of Runt-Runt cooperative interactions such as those that characterize many transcription factors [Park et al., 2019, Estrada et al., 2016b, He et al., 2010, Segal et al., 2008, Ptashne, 2004]. However, adding a Runt-Runt cooperativity term, *ω_rr_*, was insufficient to account for the observed regulatory behavior (Fig. 6C; Fig. S13 more thoroughly analyzes this discrepancy). A second alternative consists in incorporating a Runt-Runt-RNAP higher-order cooperativity term, *ω_rrp_*. While the best MCMC 1ts revealed signi1cant improvements in predictive power, important deviations still existed for the [110] construct (Fig. 6D, right; Fig. S14 more thoroughly analyzes the MCMC inference results).

Not surprisingly, given the agreement of the higher-order cooperativity model with the data for the [011] and [101] constructs (Fig. 6D, left and center), this agreement persisted when both Runt-Runt cooperativity and Runt-Runt-RNAP higher-order cooperativity were considered (Fig. 6E, left and center). However, including these two cooperativities also significantly improved the ability of model at explaining the [110] experimental data (Fig. 6E, right). Thus, while higher-order cooperativity is the main interaction necessary to quantitatively describe repression by two Runt repressors, pairwise cooperativity also needs to be invoked. This conclusion is supported by our MCMC sampling: posterior distributions for the Runt-Runt cooperativity term are not well constrained for the [011] or [101] constructs, whereas Runt-Runt-RNAP higher-order cooperativity is constrained very well across all constructs (Fig. S15D; Fig. S15 more thoroughly analyzes the MCMC inference results). As a result, accounting for both pairwise and higher order cooperativity is necessary for the model to explain the observed rate of RNAP loading of all three constructs.

The higher-order cooperativity revealed by our analysis can lead to more or less repression than predicted by the independent repression model, motivating us to determine the magnitude of this cooperativity across constructs. To make this possible, we inferred the magnitude of the Runt-Runt cooperativity *ω_rr_* and the Runt-Runt-RNAP higher-order cooperativity *ω_rrp_*. As shown in Figure 6F, depending on the spatial arrangement of Runt binding sites, the Runt-Runt-RNAP higher-order cooperativity term *ω_rrp_* can be below or above 1. Note that, in doing these 1ts, we 1rst set the Runt-Runt cooperativity, *ω_rr_*, values for [011] and [101] to 1 because, as we had demonstrated in Figure 6D, only the higher-order Runt-Runt-RNAP cooperativity was necessary. Thus, different placements of Runt molecules on the enhancer lead to distinct higher-order interactions with RNAP which, in turn, can result in less or more repression than predicted by a model where Runt molecules act independently of each other.

### 2.6 Repression by three-Runt binding sites also requires higher-order cooperativity

Building on our success in deploying thermodynamic models to explain repression by one- and two-Runt binding sites, we investigated repression by three-Runt binding sites. First, we accounted for pairwise interactions between Runt and RNAP, which were inferred from measurements of the one-Runt binding site constructs (Fig. 1B), yielding *ω_rp_*_[001]_ *, ω_rp_*_[010]_, and *ω_rp_*_[100]_ from [001], [010], and [100]. Second, we considered pairwise protein-protein interactions between Runt molecules (Fig. 1C), which were inferred from the two-Runt binding sites constructs through the parameters *ω_rr_*_[011]_ *, ω_rr_*_[101]_, and *ω_rr_*_[110]_. Finally, we incorporated Runt-Runt-RNAP higher-order cooperativity acquired from the two-Runt binding sites constructs (Fig. 1C) captured by *ω_rrp_*_[011]_ *, ω_rrp_*_[101]_, and *ω_rrp_*_[110]_. we tested our model predictions using a similar scheme to that described in the previous section: we generated a parameter-free prediction for the initial rate of transcription by using the inferred parameters from the one- and two-Runt binding sites constructs, including the pairwise and higher-order interactions described above.

Figure 7A shows the resulting parameter-free prediction. As seen in the figure, our model could not qualitatively recapitulate the experimental data as it predicted too much repression. Such disagreement suggests that additional regulatory interactions are at play. Building on the need for higher-order cooperativity in the two-Runt binding site case, we propose the existence of higher order cooperativities necessary to describe regulation by three Runt molecules—Runt-Runt-Runt higher-order cooperativity, *ω_rrr_* and Runt-Runt-Runt-RNAP higher-order cooperativity, *ω_rrrp_* (Fig. 1D). The resulting expression for the predicted rate of transcription in the presence of all these sources of cooperativity is shown in Equation S10 in Section S2. Importantly, we did not try to 1nd the optimal value for these higher-order cooprativities through 1tting. Instead, our objective was to determine whether the addition of any of these new parameters was sufficient to explain our data. When including only a Runt-Runt-Runt-RNAP higher-order cooperativity parameter of *ω_rrrp_* = 2300, our model recapitulated the experimental data (Fig. 7B). Thus, our results further support the view in which the addition of Runt repressor binding motifs in an enhancer cannot be explained by a simple additive interaction between each bound repressor. Rather, their combinatorial effect must be taken into account.

**Figure 7.**
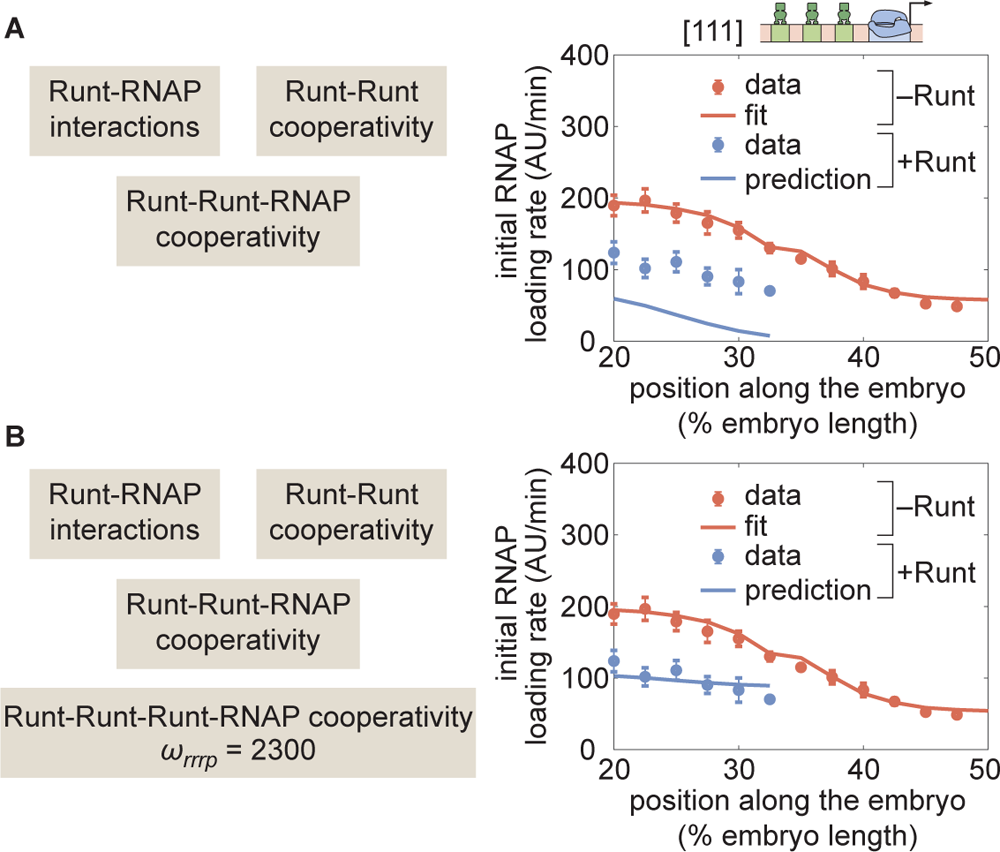
Prediction for *hunchback* P2 with three-Runt binding sites and multiple sources of cooperativity. **(A)** Prediction using previously inferred Runt-RNAP, Runt-Runt, and Runt-Runt-RNAP cooperativity parameters. **(B)** Prediction as in (A), but incorporating an additional Runt-Runt-Runt-RNAP higher-order cooperativity parameter of *ω_rrrp_* = 2300, corresponding to roughly 8 *k_B_T* of free energy. (Data points represent mean and standard error of the mean over >3 embryos.)

## 3 Discussion

One of the challenges in generating predictions to probe thermodynamic models is that, often, these models are contrasted against experimental data from endogenous regulatory regions [Segal et al., 2008, Sayal et al., 2016, Park et al., 2019]. Here, the presence of multiple binding sites for several transcription factors—known and unknown [Vincent et al., 2016]—leads to models with a combinatorial explosion of free parameters. Like the proverbial elephant that can be 1t with four parameters [Mayer et al., 2010], experiments with endogenous enhancers typically contain enough parameters to render it possible to explain away apparent disagreement between theory and experiment [Garcia et al., 2020].

To close this gap, synthetic minimal enhancers have emerged as an attractive alternative to endogenous enhancers [Fakhouri et al., 2010, Sayal et al., 2016, Park et al., 2019, Crocker et al., 2016]. Here, the presence of only a handful of transcription factor binding sites and the ability to systematically control their placement and affinity dramatically reduce the number of free parameters in the model [Garcia et al., 2020]. Inferences performed on these synthetic constructs could then inform model parameters that would make it possible to quantitatively predict transcriptional output of *de novo* enhancers [Sayal et al., 2016].

Building on these works, in the present investigation we sought to predict how the Runt repressor, which counteracts activation by Bicoid along the anterior-posterior axis of the early fly embryo [Chen et al., 2012], dictates the output level of transcription. To dissect repression, a strong and detectable level of expression in the absence of the repressor was needed, prompting us to choose a simple system of synthetic enhancers based on the strong *hunchback* P2 minimal enhancer [Garcia et al., 2013, Chen et al., 2012]. This enhancer has been carefully dissected in terms of its activator Bicoid and the pioneer-like transcription factor Zelda in the early embryo [Driever and Nusslein-Volhard, 1988, Garcia et al., 2013, Park et al., 2019, Eck et al., 2020], making it easier to identify neutral sequences within the enhancer for introducing Runt binding sites [Chen et al., 2012]. Further, when inserted into *hunchback* P2, Runt binding site number determines the level of transcription incrementally [Chen et al., 2012]. Thus, *hunchback* P2 provided an ideal scaffold onto which to quantitatively and systematically dissect repression by Runt.

Previous studies using synthetic enhancers relied on measurements of input transcription factor patterns using fluorescence immunostaining, and of cytoplasmic mRNA patterns using fluorescence in situ hybridization (FISH) or single-molecule FISH. These fixed-tissue techniques have key differences from the live-imaging approach adopted here. First, given the dynamical nature of development, it is necessary to know when data were acquired. Doing so with high temporal resolution using FISH is challenging, although it can be accomplished to some degree by synchronizing embryo deposition before fixation [Park et al., 2019]. Second, while most transcription factors directly dictate the rate of RNAP loading, and hence the rate of mRNA production [Spitz and Furlong, 2012, Garcia et al., 2013, Eck et al., 2020], typical FISH measurements report on the accumulated mRNA in the cytoplasm, which is a convolution of all processes of the transcription cycle—initiation, elongation, and termination [Liu et al., 2021, Alberts, 2015]—as well as mRNA nuclear export dynamics, diffusion, and degradation. These processes could be modulated in space and time, potentially confounding measurements. Here, we overcame these challenges by using the MS2 technique to precisely time our embryos and acquire the rate of transcription initiation.

Interestingly, our initial dissection of constructs containing various combinations of Runt binding sites, but in the absence of Runt protein, revealed that unrepressed gene expression levels depend strongly on the number and placement of the binding sites within the enhancer (Fig. 4A). These results challenge previous assumptions that unregulated gene expression levels would stay unchanged as enhancer architecture is modulated [Sayal et al., 2016, Fakhouri et al., 2010, Barr et al., 2017], but they are in accordance with observations in bacterial systems [Garcia et al., 2012]. As a result, our measurements call for accounting for unregulated levels in future quantitative dissections of eukaryotic enhancers, or to study relative magnitudes such as the fold-change in gene expression that has driven the dissection of bacterial transcriptional regulation [Phillips et al., 2019].

Once we accounted for this difference in unrepressed gene expression levels, we determined that the repression profiles obtained for constructs bearing one-Runt binding site could be described by a simple thermodynamic model (Fig. 2). Specifically, we showed that the same dissociation constant described Runt binding regardless of the position of its binding site along the enhancer (Fig. 5A). Further, the Runt-RNAP interaction terms describing repressor action decreased as the binding site was placed farther from the promoter (Fig. 5C), qualitatively consistent with a “direct repression” model in which Runt needs to physically contact RNAP in order to realize its function [Jaynes and O’Farrell, 1991, Gray et al., 1994, Hewitt et al., 1999].

Although our model recapitulated repression by a one-Runt binding site, the inferred parameters were insufficient to quantitatively predict repression by two-Runt binding sites (Fig. S6B). These results suggest that multiple repressors do not act independently of each other. Instead, new parameters describing both Runt-Runt cooperativity and Runt-Runt-RNAP higher-order cooperativity had to be incorporated into our models to quantitatively describe Runt action in these constructs (Fig. S6C-E).

While we have long known about protein-protein cooperative interactions [Ackers et al., 1982], in the last few years it has become clear that higher-order cooperativity can also be at play in eukaryotic systems [Estrada et al., 2016a, Park et al., 2019, Biddle et al., 2020] as well as in bacteria [Dodd et al., 2004] and archaea [Peeters et al., 2013]. The existence of this higher-order cooperativity suggests that, to predict gene expression from DNA sequence, it might be necessary to build an understanding of the many simultaneous interactions that precede transcriptional initiation. Our discovery of higher-order cooperativity in the action of multiple Runt molecules opens up new avenues to uncover the molecular nature of this phenomenon. For example, following an approach developed in [Park et al., 2019], it could be possible to determine whether and how these cooperativity parameters are modulated upon perturbation of molecular players such as the Groucho or CtBP co-repressors, Big-brother, a co-factor facilitating the Runt binding to DNA, and components of the mediator complex [Park et al., 2019, Courey and Jia, 2001, Walrad et al., 2011]. Indeed, [Park et al., 2019] recently showed that co-activators and mediator units are involved in dictating the magnitude of similar higher-order cooperativity terms in activation by Bicoid. Thus, our thermodynamic models provide a lens through which to dissect the molecular underpinnings of Runt interactions with itself and with the transcriptional machinery.

Notably, the need to invoke cooperative interactions as more Runt binding sites are being added opposes our goal of predicting complex regulatory architectures from experiments with simpler architectures without the need to invoke new parameters. However, it will be interesting to determine whether more parameters need to be invoked as the number of Runt binding sites increases beyond three, or whether the parameters already inferred are sufficient to endow our models with parameter-free predictive power.

Importantly, while our model adopted a “direct repression” view of the mechanism of Runt action, other mechanisms of repression such as “quenching” could also describe the data. While all such models call for higher-order cooperativity to describe the data (Supplementary Section S5), our data cannot differentiate among those models. Thus, we did not attempt to distinguish different molecular mechanisms of Runt transcriptional repression.

Finally, even though the work presented here has relied exclusively on thermodynamic models, it is important to note that a much more general approach based on non-equilibrium models could also be appropriate for describing our data. Indeed, an increasing body of work over the last few years has provided evidence for the necessity of invoking these more complex models in the context of transcriptional regulation in eukaryotes [Estrada et al., 2016a, Li et al., 2018, Park et al., 2019, Eck et al., 2020]. In future work, it will be interesting to determine whether, when our data is viewed through the lens of these non-equilibrium models, invoking higher-order cooperativity is still necessary or whether, instead, simple pairwise protein-protein interactions suffice to reach an agreement between theory and experiment.

Overall, the work presented here establishes a framework for systematically and quantitatively studying repression in the early fly embryo. As showcased here, synthetic enhancers based on the *hunchback* P2 minimal enhancer constitute an ideal scaffold for the study of other repressors in early fly embryos. For example, we envision that this approach could be used to dissect repression by other transcription factors such as *Capicua* or *Krüppel* [Löhr et al., 2009, Sauer and Jackle, 1991, Papagianni et al., 2018, Chen et al., 2012], and to probe observations of multiple repressors working together to oppose activation by Bicoid in establishing gene expression patterns along the anterior-posterior axis [Chen et al., 2012, Briscoe and Small, 2015]. We anticipate that a similar approach could be used to dissect repression along the dorso-ventral axis of the embryo, by for example, adding repressor binding sites to well-established reporter constructs that are only regulated by the Dorsal activator [Jiang and Levine, 1993]. Critically, we need to understand not only how one species of repressor works in concert with an activator, but also how multiple species of repressors work together as a system. The approach presented here provides a way forward for predictively understanding the complex gene regulatory network that shapes gene expression patterns in the early fly embryo.

## 4 Materials and Methods

### 4.1 Generation of synthetic enhancers with MS2 reporters

The synthetic enhancer constructs used in this study are based off of Chen et al. [2012]. In summary, the *hunchback* P2 enhancer was used as a scaffold to introduce Runt binding sites at different positions that are thought to be neutral (i.e. these Runt binding sites do not interfere with any other obvious binding sites for other transcription factors in the early *Drosophila* embryos as shown in Fig. S9). For the three positions chosen to introduce Runt binding sites in Chen et al. [2012], the Gene Synthesis service from Genscript was used to generate synthetic enhancers with all possible configurations of zero-, one-, two-, and three-Runt binding sites in *hunchback* P2 as shown in Figure 1A. The enhancer sequences were placed into the original plasmid pIB backbone [Chen et al., 2012] using the Gene Fragment Synthesis service in Genscript, followed by the *even-skipped* promoter, and 24 repeats of MS2v5 loops [Wu et al., 2015], the *lacZ* coding sequence, and the *a*-Tubulin 3’UTR sequence [Chen et al., 2012]. These plasmids were injected into the 38F1 landing site using the RMCE method [Bateman et al., 2006] by BestGene Inc. Flies were screened by selecting for white eye color and made homozygous. The orientation of the insertion was determined by genomic PCR to ensure a consistent orientation across all of our constructs. Specifically, we used two sets of primers that each amplified one of these two possible orientations: “Upward”, where the forward primer binds to a genomic location outside of 38F1 (TTCTAGTTCCAGTGAAATCCAAGCA) and the reverse primer binds to a location in our reporter transgene (ACGCCAGGGTTTTCCCAG), and “Downward”, where the forward primer remains the same as the “Upward” set and the reverse primer binds to a location in our reporter transgene (CTCTGTTCTCGCTATTATTCCAACC) when the insertion is the opposite orientation to the “Upward” orientation. As a result, only amplicons from either one of the orientations of insertion in the 38F1 landing site can be obtained. We chose the “Downward” orientation for all our constructs.

### 4.2 CRISPR-Cas9 knock-in of the green LlamaTag in the endogenous *runt* locus

We used CRISPR-Cas9 mediated Homology Directed Repair (HDR) to insert the LlamaTag against eGFP into the N-terminal of the *runt* endogenous locus [Bothma et al., 2018, Gratz et al., 2015]. The donor plasmid was constructed by stitching individual fragments—PCR amplified left/right homology arms from the endogenous *runt* locus roughly 1 kb in length each, LlamaTag, and pHD-scarless vector—using Gibson assembly [Gratz et al., 2015]. The PAM sites in the donor plasmid were mutated such that the Cas9 only cleaved the endogenous loci, not the donor plasmid, without changing the amino acid sequence of the Runt protein. The final donor plasmid contained the 3xP3-dsRed marker such that dsRed is expressed in the fly eye and ocelli for screening. Positive transformant flies were screened using a fluorescence dissection scope and set up for single fly crosses to establish individual lines that were then veri1ed with PCR ampli1cation and Sanger sequencing (UC Berkeley Sequencing Facility). Importantly, this *llamaTag-runt* allele rescues development to adulthood as a homozygous. Thus we concluded that the LlamaTag-Runt allele can be used to monitor the behavior of endogenous Runt protein.

### 4.3 Fly strains

Transcription from the synthetic enhancer reporter constructs was measured by using embryos from crossing *yw;his2av-mRFP1;MCP-eGFP(2)* females and *yw;synthetic enhancer-MS2v5-lacZ;+* males as described in [Garcia et al., 2013, Eck et al., 2020, Lammers et al., 2020]. eGFP-Bicoid measurements were performed using the fly line from [Gregor et al., 2007]. The LlamaTag-Runt measurements were done using the fly line *LlamaTag-Runt; +; vasa-eGFP, His2Av-iRFP* illustrated in Table 2. Briefly, eGFP was supplied by a *vasa* maternal driver. Females carrying both the LlamaTag-Runt and the *vasa*-driven eGFP were crossed with males carrying the LlamaTag-Runt, the progeny from this cross were imaged and then recovered to determine the embryo’s sex using PCR. PCR was run with three sets of primers: Y chr1 (Forward: CGATCCAGCCCAATCTCTCATAT-CACTA, Reverse: ATCGTCGGTAATGTGTCCTCCGTAATTT), Y chr2 (Forward: AACGTAACCTAGTCGGATTG-CAAATGGT, Reverse: GAGGCGTACAATTTCCTTTCTCATGTCA), and Auto1 (Forward: GATTCGATGCA-CACTCACATTCTTCTCC, Reverse: GCTCAGCGCGAAACTAACATGAAAAACT). Two of primer sets (Y chr1 and Y chr2) bind to the Y chromosome while the other one (Auto1) binds to the autosome and constitutes a positive control [Lott et al., 2011].

To generate the embryos that are zygotic null for the *runt* allele, we used a fly cross scheme consisting of two crosses. In the first generation, we crossed *LlamaTag-Runt;+;+* males with *run3/FM6;+;MCP-eGFP(4F),his2av-mRFP1* females. *run3* is the null allele for *runt*, missing around 5 kb including the coding sequence of the *runt* locus [Gergen and Butler, 1988, Chen et al., 2012]. The *MCP-eGFP(4F)* transgene expresses approximately twice the amount of MCP protein than the *MCP-eGFP(2)* [Garcia et al., 2013, Eck et al., 2020] and thus results in similar levels of MCP to those of *MCP-eGFP(2)* in the trans-heterozygotes. The female progeny from this cross, *LlamaTag-Runt/run3;+;MCP-eGFP(4F),his2av-mRFP1/+* was then crossed with males whose genotype was *LlamaTag-Runt/Y;synthetic enhancer-MS2v5-lacZ;+* to produce the embryos that we used for live imaging. The resulting embryos carried maternally supplied MCP-eGFP and His-RFP for visualization of nascent transcripts and nuclei. The X chromosome contained LlamaTag-Runt allele or *run3* null allele. We could differentiate between these two genotypes because, when the embryo had the Runt allele, a stripe pattern would appear in late nc14. We imaged all embryos until late nc14 to make sure that we were capturing the nulls.

### 4.4 Sample preparation and data collection

Sample preparation was done following the protocols described in Garcia et al. [2013]. Briefly, embryos were collected, dechorionated with bleach for 1-2 minutes, and then mounted between a semipermeable membrane (Lumox 1lm, Starstedt, Germany) and a coverslip while embedded in Halocarbon 27 oil (Sigma-Aldrich). Live imaging was performed using a Leica SP8 scanning confocal microscope, a White Light Laser and HyD dectectors (Leica Microsystems, Biberach, Germany). Imaging settings for the MS2 experiments with the presence of MCP-eGFP and Histone-RFP were the same as in Eck et al. [2020] except that we used 1024×245 (pixels) format to image a wider field of view along the anterior-posterior axis. The settings for the eGFP-Bicoid measurements were the same as described in Eck et al. [2020].

The settings for the eGFP:LlamaTag-Runt measurements were similar to that of eGFP-Bicoid except for the following. To increase our imaging throughput, we utilized the “Mark and Position” functionality in the LASX software (Leica SP8) to image 5-6 embryos simultaneously. To account for the decreased time resolution, we lowered the z-stack size from 10 *µ*m to 2.5 *µ*m, keeping the 0.5 *µ*m z-step. By doing this, we could maintain 1 minute frame rate for each imaged embryo. Additionally, these flies expressed Histone-iRFP, instead of Histone-RFP as in Eck et al. [2020], so that we used a 670 nm laser at 40 *µ*W (measured at a 10x objective) for excitation of the histone channel, and the HyD detector was set to a 680 nm-800 nm spectral window.

### 4.5 Image Analysis

Images were analyzed using custom-written software (MATLAB, mRNA Dynamics Github repository) following the protocol in Garcia et al. [2013] and Eck et al. [2020]. Briefly, this procedure involved segmentation and tracking of nuclei and transcription spots. First, segmentation and tracking of individual nuclei were done using the histone channel as a nuclear mask. Second, segmentation of each transcription spot was done based on its fluorescence intensity and existence over multiple z-stacks. The intensity of each MCP-GFP transcriptional spot was calculated by integrating pixel intensity values in a small window around the spot and subtracting the background fluorescence measured outside of the active transcriptional locus. When there was no detectable transcriptional activity, we assigned NaN values for the intensity. The tracking of transcriptional spots was done by using the nuclear tracking and proximity of transcriptional spots between consecutive time points. The nuclear protein fluorescence intensities from the eGFP-Bicoid and LlamaTag-Runt fly lines, which we use as a proxy for the protein nuclear concentration, were calculated as follows. Using the nuclear mask generated from the histone channel, we performed the same nuclear segmentation and tracking as described above for the MS2 spots. Then,for every z-section, we extracted the integrated fluorescence over a 2*µm* diameter circle on the xy-plane centered on each nucleus. For each nucleus, the recorded fluorescence corresponded to the z-position where the fluorescence was maximal. This resulted in an average nuclear concentration as a function of time for each single nucleus. These concentrations from individual nuclei were then averaged over a narrow spatial window (2.5% of the embryo length) to generate the spatially averaged protein concentration reported in the main text. For the eGFP:LlamaTag-Runt datasets, we had to subtract the background eGFP fluorescence due to the presence of an unbound eGFP population [Bothma et al., 2018]. We used the same protocol described in Bothma et al. [2018] and in the Supplementary Section S7 to extract this background.

### 4.6 Bayesian inference procedure: Markov Chain Monte Carlo sampling

Parameter inference was done using the Markov Chain Monte Carlo (MCMC) method. We used a well-established package *MCMCstat* that uses an adaptive MCMC algorithm [Haario et al., 2006, 2001]. A detailed description on how we performed the MCMC parameter inference, for example setting the priors and bounds for parameters, is illustrated in Supplementary Section S4.

### 4.7 Biological Materials

**Table 1.** List of plasmids used to create the transgenic fly lines used in this study.

| Plasmids |  |
| --- | --- |
| Name (hyperlinked to Benchling) | Function |
| pIB-hbP2-evePr-MS2v5-LacZ-Tub3UTR | [000]-MS2v5 reporter construct |
| pIB-hbP2+r1-far-evePr-MS2v5-LacZ-Tub3UTR | [100]-MS2v5 reporter construct |
| pIB-hbP2+r1-mid-evePr-MS2v5-LacZ-Tub3UTR | [010]-MS2v5 reporter construct |
| pIB-hbP2+r1-close-evePr-MS2v5-LacZ-Tub3UTR | [001]-MS2v5 reporter construct |
| pIB-hbP2+r2-2+3-evePr-MS2v5-LacZ-Tub3UTR | [011]-MS2v5 reporter construct |
| pIB-hbP2+r2-1+3-evePr-MS2v5-LacZ-Tub3UTR | [101]-MS2v5 reporter construct |
| pIB-hbP2+r2-1+2-evePr-MS2v5-LacZ-Tub3UTR | [110]-MS2v5 reporter construct |
| pIB-hbP2+r3-evePr-MS2v5-LacZ-Tub3UTR | [111]-MS2v5 reporter construct |
| pHD-scarless-LlamaTag-Runt | Donor plasmid for LlamaTag-Runt CRISPR knock-in fusion for the N-terminal |
| pU6:3-gRNA(Runt-N-2) | gRNA plasmid for LlamaTag-Runt CRISPR knock-in fusion for the N-terminal |
| pCasper-vasa-eGFP | *vasa* maternal driver for ubiquitous eGFP expression in the early embryo |

**Table 2.** List of fly lines used in this study and their experimental usage

| Fly lines |  |
| --- | --- |
| Genotype | Use |
| *LlamaTag-Runt*; +; *vasa-eGFP*, *His2Av-iRFP* | Visualize LlamaTagged Runt protein and label nuclei |
| *LlamaTag-Runt*; +; *MCP-eGFP(4F)*, *His2Av-iRFP* | Visualize LlamaTagged Runt protein, nascent transcripts and label nuclei |
| *run3/FM6*; +; + | Visualize LlamaTagged Runt protein, nascent transcripts and label nuclei |
| *yw*; *His2Av-mRFP*; *MCP-eGFP* | Females to label nascent RNA and nuclei |
| *yw*; [000]-*MS2v5* ; + | Males carrying the MS2 reporter transgene |
| *yw*; [100]-*MS2v5* ; + | Males carrying the MS2 reporter transgene |
| *yw*; [010]-*MS2v5* ; + | Males carrying the MS2 reporter transgene |
| *yw*; [001]-*MS2v5* ; + | Males carrying the MS2 reporter transgene |
| *yw*; [011]-*MS2v5* ; + | Males carrying the MS2 reporter transgene |
| *yw*; [101]-*MS2v5* ; + | Males carrying the MS2 reporter transgene |
| *yw*; [110]-*MS2v5* ; + | Males carrying the MS2 reporter transgene |
| *yw*; [111]-*MS2v5* ; + | Males carrying the MS2 reporter transgene |

## Acknowledgements

We are grateful to Armando Reimer, Brandon Schlomann, Elizabeth Eck, Gabriella Martini, Jacques Bothma, Jeehae Park, Jia Ling, Matthew Norstad, Michael Eisen, Nicholas Lammers, Nipam Patel, Rob Phillips, Simon Alamos, Xavier Darzacq,and Yasemin Kirişçioğlu for their guidance and comments on our manuscript. This work was supported by the Burroughs Wellcome Fund Career Award at the Scientific Interface, the Sloan Research Foundation, the Human Frontiers Science Program, the Searle Scholars Program, the Shurl and Kay Curci Foundation, the Hellman Foundation, the NIH Director’s New Innovator Award (DP2 OD024541-01), and NSF CAREER Award (1652236) to HGG, and a KFAS scholarship to YJK.

## S1 Derivation of the general thermodynamic model for the *hunchback* P2 enhancer

In this section, we rederive the thermodynamic model presented in the main text, now without the assumption of strong Bicoid-Bicoid cooperativity. The equilibrium thermodynamic modeling framework that we used in this paper is described in more detail in Bintu et al. [2005b,a].

We start by modeling the case of *hunchback* P2 without any Runt binding sites, which is believed to have at least six Bicoid binding sites [Park et al., 2019, Driever et al., 1989]. As shown by the states and weights presented in Figure S1A, in our thermodynamic model, we assume that the six Bicoid binding sites have the same dissociation constant given by *K_b_*, and we posit that RNAP-promoter binding is governed by a dissociation constant given by *K_p_*. We also assume pairwise cooperativity between Bicoid molecules given by *ω_b_*, and cooperativity between each Bicoid molecule and RNAP given by *ω_bp_*. For simplicity, we will use the dimensionless parameters *b* = [*Bicoid*]/*K_b_* and *p* = [*RNAP*]/*K_p_*, where [*Bicoid*], and [*RNAP*] are the concentrations of Bicoid and RNAP, respectively, and *K_b_* and *K_p_* are their corresponding dissociation constants.

We factor the total partition function into two categories: *z_b_* corresponding to states that only have Bicoid bound, and *z_bp_* describing states with both Bicoid and RNAP bound. Then then calculate each component separately. The sum of microstates for *z_b_* is

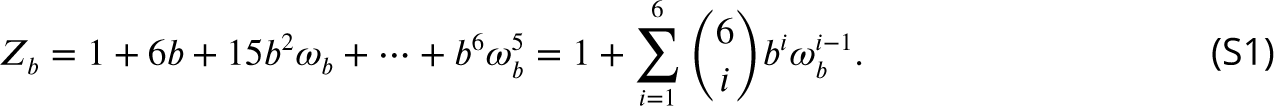

Using the binomial theorem, we can simplify Equation S1 leading to

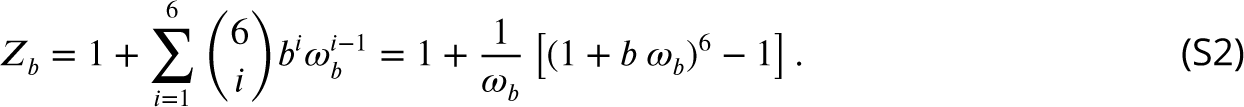

Using the same logic, we obtain *z_bp_* such that

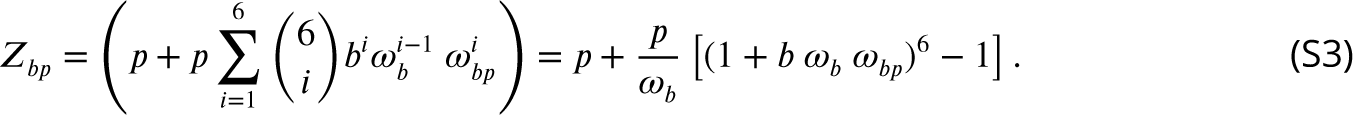

Using these two partition functions, we then calculate the probability of the promoter being bound by RNAP, *p_bound_* as

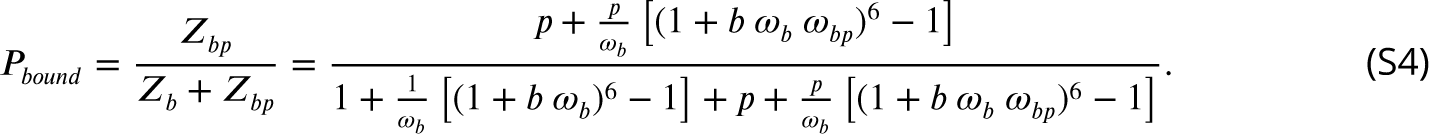

Following recent work [Gregor et al., 2007, Park et al., 2019], we now assume that the Bicoid-Bicoid pairwise cooperativity is very strong (*ω_b_ »* 1). We can then simplify Equation S4 to obtain

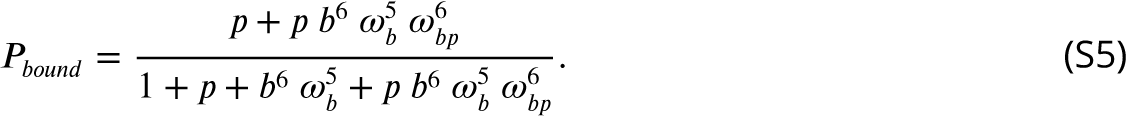

If we now define a new binding constant for Bicoid, 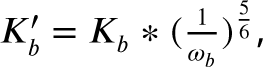, such that *b*^1^ = *b ω*_b_^5/6^, and a new cooperativity term between Bicoid and RNAP given by *ω*_bp_^1^ = *ω*_bp’_^6^, we can then rewrite Equation S5 as

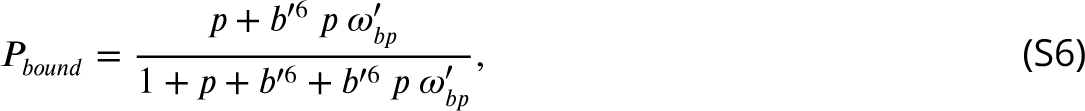

which is the expression we use throughout the main text. Thus, strong pairwise cooperativity between Bicoid molecules leads to a functional form where only the state with all Bicoid molecules bound remain (six in this case). This strong cooperativity can explain the sharp step-like expression pattern along the embryo’s anterior-posterior axis of the *hunchback* gene (Fig. 3J; Gregor et al. [2007], Park et al. [2019], Driever and Nusslein-Volhard [1988, 1989]).

## S2 Derivation of the general and simpler thermodynamic model for the *hunchback* P2 enhancer with one Runt binding site

Having derived the equation for the strong cooperative binding of Bicoid to the wild-type *hunchback* P2 enhancer, we will now extend that model to the case of *hunchback* P2 with one Runt binding site. The corresponding states and weights of our full model are shown in Figure S2A.

Using a similar logic for calculating the partition functions as described in the previous section, we can compute the probability of the promoter being bound by RNAP as

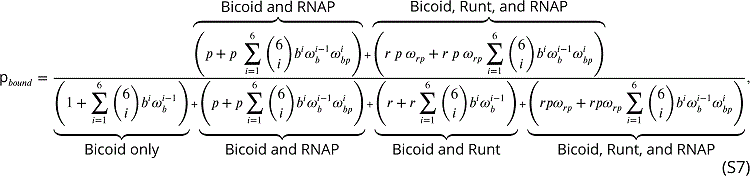

where, in addition to the parameters defined in the above section for the wild-type *hunchback* P2 case in the absence of Runt, we have added two parameters: the dissociation constant for Runt given by *K_r_*, and a Runt-RNAP interaction term (an anti-cooperativity), *ω_rp_*. Using the binomial theorem as in Equation S2, we can simplify Equation S7 to obtain

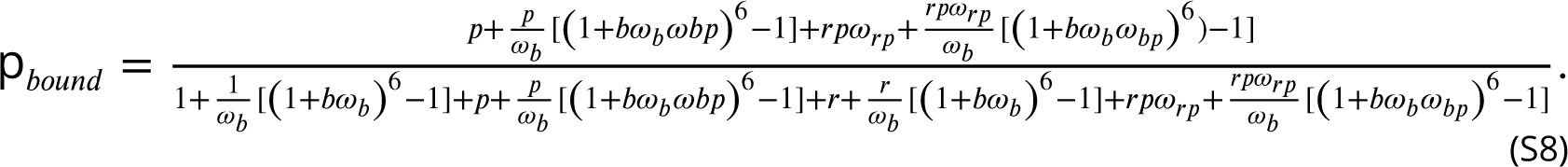

We now again assume that Bicoid-Bicoid cooperativity is very strong such that *ω_b_ »* 1. Then, we can combine Equation S8 with Equation 1 to obtain

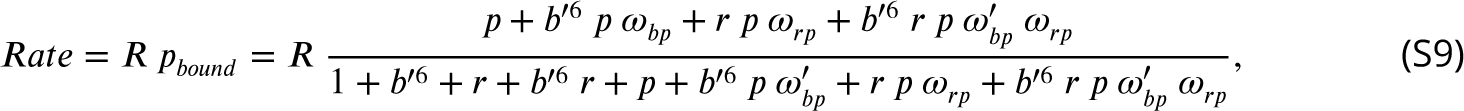

where the new parameters, *b*^1^ and *w*^1^ are de1ned in the same way as in Equation S6. The effective states and weights remaining after taking this limit are shown in Figure S2B. Similarly, we can derive expressions for *p_bound_* in the presence of two and three Runt binding sites, and in the strong Bicoid-Bicoid cooperativity limit in order to obtain the predictions used throughout this text. We show this expression for two Runt binding sites in Equation 5. Further, for the case of repression by three Runt binding sites, the rate of transcription is given by

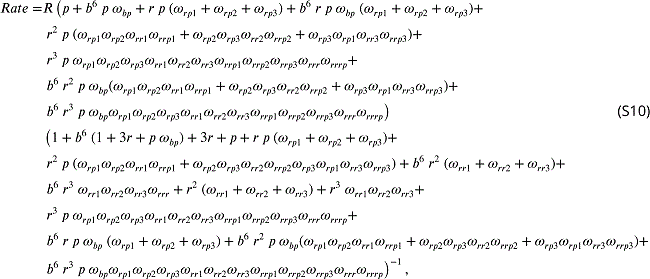

where the parameters are defined as in Figure 1 and Section 2.6.

## S3 Comparing using static versus dynamic transcription factor concentrations as model inputs

In this section, we tested whether using static, time-averaged transcription factor concentration profiles yielded comparable theoretical predictions than when instead acknowledging the fact that input transcription factor concentration changes over time. Briefly, we compared the predicted rate of transcription calculated in two ways: (1) time-averaging the instantaneous rate from the dynamic transcription factor concentration pro1les over a speci1ed time window (from 5 to 10 minutes from the 13th anaphase) and (2) using static input transcription factors already time-averaged over the same time window.

As a concrete example, we focused on the *hunchback* P2 enhancer with one Runt binding site. We calculated the predicted rate of transcription using the thermodynamic model given by Equation 2. First, we performed this calculation using the dynamic concentration profiles of Bicoid and Runt shown in Figure 3B and D, respectively. Briefly, the terns *b* and *r* in Equation 2 now become functions of time such that

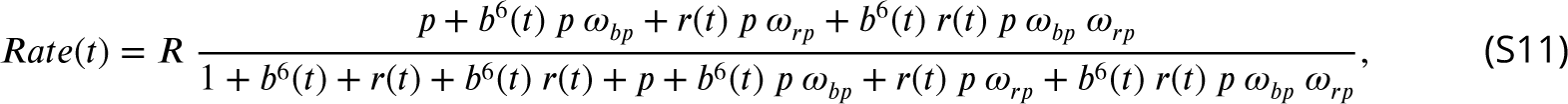

where *b*(*t*) = [*Bicoid*](*t*)/*K_b_* and *r*(*t*) = [*Runt*](*t*)/*K_r_*. We choose a set of reasonable values for the model parameters to illustrate the calculation of *Rate*(*t*) at 30% of the embryo length. The resulting dynamic rate of transcription profile is shown in Figure S3A (blue curve). We then use this profile to calculate the time-averaged rate of transcription over the time window of 5 to 10 minutes from the 13th anaphase, resulting in the green area shown in Figure S3A.

The predicted average rate of RNAP loading given dynamic input transcription factors can be compared to the predicted rate of RNAP loading given the average input concentrations that we used throughout the main text (Fig. 3E). Specifically, we plug the static concentration profiles of Bicoid and Runt shown in Figure 3E into Equation 2 to obtain the red area shown in Figure S3A. As shown in the figure, the predicted rate of transcription obtained by these two analysis methodologies are equivalent within error.

Finally, we performed this comparison between different approaches to calcualte the rate of transcription as a function of position along the embryo (from 20% to 70% of the embryo length). As shown in Figure S3B, the resulting spatial profiles are comparable within error. Thus, we have shown that our approach of using time-averaged, static transcription factor concentrations as inputs to our model yield quantitatively equivalent result as accounting for the dynamic concentration profiles of these transcription factors.

## S4 Markov Chain Monte Carlo inference protocol

Markov Chain Monte Carlo (MCMC) sampling is a widely used technique for robust parameter estimation using Bayesian statistics [Geyer and Thompson, 1992, Sivia and Skilling, 2006]. We used the MATLAB package *MCMCstat*, an adaptive MCMC technique, which we could directly implement downstream of our data analysis pipeline [Haario et al., 2006, 2001]. Detailed instructions on how to implement the *MCMCstat* package can be found in https://mjlaine.github.io/mcmcstat/.

MCMC allows for an estimation of the set of parameter values of a model that best explain the experimental data along with their associated errors. In this work, we used MCMC to infer the set of best 1t values of the parameters in our thermodynamic models given the observed profile of the rate of transcription initiation along the anterior-posterior axis of the embryo.

MCMC calculates a Bayesian posterior probability distribution of each free parameter given the data by stochastically sampling different parameter values. For a given set of observations *D* and a model with parameters *θ*, the posterior probability distribution of a particular set of values is given by Bayes’ theorem

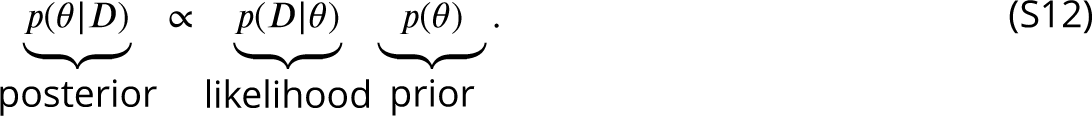

The prior function represents the *a priori* assumption about the probability distribution of parameter values *θ*. Here, we assumed a uniform prior distribution for all parameters to reflect our ignorance about the model parameters within the following intervals:

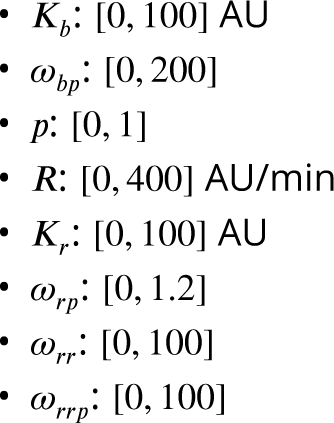

These intervals were justified using the following arguments.

First, because we observed a gradual modulation of the rate of transcription by both Bicoid and Runt in the middle region of the embryo we reasoned that the binding sites for these transcription factors were not saturated. As a result, we posited that the real dissociation constant should be between the minimum and maximum measured values of Bicoid and Runt (Fig. S10). Our measurements of Bicoid and Runt concentration yield fluorescence values over the 0-100 AU range for the embryo region that we used for contrasting our model and experimental data (20-50% of the embryo length), such that the dissociation constants (*K_b_* and *K_r_*) should not exceed the maximum value of the Bicoid or Runt concentration.

Second, *ω_bp_* represents the cooperativity between Bicoid complex and RNAP. In the statistical mechanics framework, this cooperativity can be expressed using the interaction energy between Bicoid and RNAP, Δε_bp_, such that *ω_bp_ = exp(−βΔε_bp_*, where 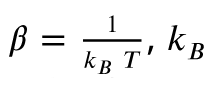 is the Boltzmann constant *k T* and *T* is the temperature. There is not much known about *in vivo* interaction energies between Bicoid and RNAP complex, thus we tried several different bounds until we found a narrow enough parameter bound with unimodal distribution of the posterior chain. As we could see from the corner plots in Figure 4C, there is a positive correlation between *K_b_* and *ω_bp_*. Thus, we constrained the *ω_bp_* intervals by 1nding an interval that gives both well-constrained *K_b_* and *ω_bp_* (Fig. 4C).

Third, *R* represents the rate of RNAP loading when the promoter is occupied, thus it is constrained by the maximum observed rate of RNAP loading (Fig. S10).

Fourth, *p* = [*RNAP*]/*K_p_*) represents the concentration of RNAP divided by its dissociation constant. Recall that the predicted rate of transcription from *hunchback* P2 in the limit where the Bicoid concentration reaches zero is given by

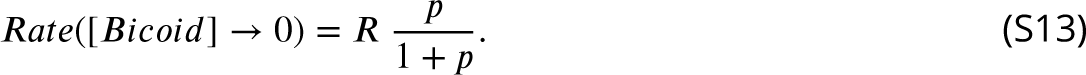

This rate of transcription at the posterior region, where Bicoid reaches zero, is much lower than that at the anterior region where Bicoid saturates given by *R* (Fig. S10). As a result, we can write the inequality

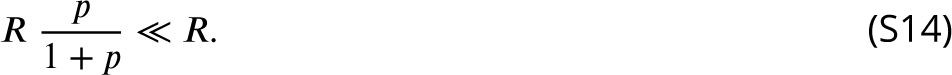

such that

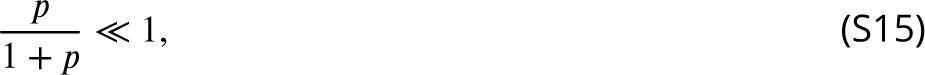

which holds if p ≪ 1.

Finally, we did not have good estimates for the intervals of either Runt-Runt cooperativity, *ω_rr_*, or higher-order cooperativity, *ω_rrp_*. Thus, we initially started with an interval of [0, 100], of the same order as the interval we used *ω_bp_*. We then explored whether this parameter bound was sufficient to give us constrained values of *ω_rr_* and *ω_rrp_*. As we showed in Figure S15D, this interval gives reasonably constrained values of *ω_rr_* and *ω_rrp_*. As shown in Figure 6 and Figure S15, we posit that the *ω_rr_* parameter is not well-constrained not because of its width of the interval, but because it is not as essential for the model 1t to the data as it is to include *ω_rrp_* into the model. Overall, our MCMC inference results as well as the corner plots shown demonstrate that our parameter intervals chosen were reasonable.

## S5 Comparison of different modes of repression

Transcriptional repressors have been classified into two broad categories: short-range and longrange, depending on the genomic length scale that they act on [Courey and Jia, 2001, Li and Gilmour, 2011]. Long-range repression is realized by the recruitment of chromatin modifiers. In contrast, short-range repressors act within 100-150 bp by interacting with nearby transcription factors or with the promoter [Li and Gilmour, 2011]. Traditionally, the molecular mechanism of short-range repressors, such as Runt, have been further classified into three categories: “direct repression”, “competition”, and “quenching” [Gray et al., 1994, Jaynes and O’Farrell, 1991, Arnosti et al., 1996, Kulkarni and Arnosti, 2005]. In “direct repression”, the repressor inhibits the binding of RNAP to the promoter (Fig. S4A). “Competition” denotes a repressor that competes with an activator for the same DNA binding location (Fig. S4B). This molecular mechanisms has been proposed for the action of Giant and Krüppel repressors on the *even-skipped* stripe 2 enhancer, where some activator and repressor binding sites partially overlap [Small et al., 1992]. Lastly, “quenching” corresponds to the case where the repressor and activator do not interact with each other directly. Instead, the repressor inhibits the activators’ action of recruiting the RNAP (Fig. S4C).

Despite several classic studies of the molecular mechanism of repressors in the early fly embryo [Gray et al., 1994, Ip et al., 1992, Bothma et al., 2011, Jaynes and O’Farrell, 1991], the mechanisms of many repressors remain unknown. Note that, even for the same repressor, the mode of repression might not be the same depending on, for example, its sequence context [Koromila and Stathopoulos, 2019, Hang and Gergen, 2017]. For example, it has been proposed that Runt repressor acts with different mechanisms in different regulatory elements of the *sloppy-paired* gene [Hang and Gergen, 2017]. In this section, we derive a thermodynamic model from each mode of repression and compare their explanatory power in the context of our data stemming from the *hunchback* P2 enhancer containing one Runt binding site. Note that, in the main text, we already developed a thermodynamic model for the “direct repression” scenario (Section S2). As a result, in this section, we focus on deriving the thermodynamic models for the “competition” and “quenching” scenarios, but repeat the result of the derivation for the “direct repression” here for ease comparison between different models.

## S5.1 Derivation of models for each scenario of repression for *hunchback* P2 with one Runt binding site

### S5.1.1 Modeling repression for *hunchback* P2 with one Runt binding site: direct repression

For completeness, we repeat the expression for the direct repression scenario as shown in Section S2 and Figure S4A. The probability of finding RNAP bound to the promoter, *p_bound_*, is calculated by dividing the sum of all statistical weights featuring RNAP by the sum of the weights of all possible microstates. The calculation of *p_bound_*, combined with Equation 1, leads to the expression

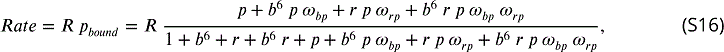

where the parameters are as defined in Figure 2.

### S5.1.2 Modeling repression for *hunchback* P2 with one Runt binding site: competition

In the competition scenario, Runt binding makes Bicoid binding less likely. This mechanism can be captured by an interaction term between Bicoid and Runt given by *ω_br_*. Building on our assumption of strong Bicoid-Bicoid cooperativity, we posit that Runt disfavors the state with six bound Bicoid molecules. We can enumerate the states and weights from Fig. S4B to calculate the *Rate* ((X *p_bound_*), which leads to

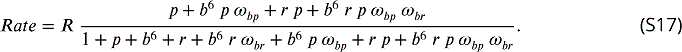

### S5.1.3 Modeling repression for *hunchback* P2 with one Runt binding site: quenching

In the quenching scenario, Runt reduces the magnitude of the cooperativity between the Bicoid complex and RNAP by a factor *ω_brp_*. We can enumerate the states and weights from Fig. S4C, leading to a rate of transcription given by

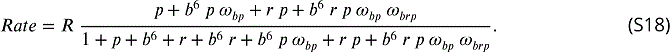

With these expressions for each repression mechanism in hand, we can now compare how each model fares against our experimental data.

## S5.2 Comparing the three models of repression with the one-Runt binding site data

We used the MCMC sampling to 1t each model to our experimentally measured initial rate of transcription over the anterior-posterior axis of the embryo. As shown in Figure S5A, B, and C, we see that all three models can explain the [100] and [010] construct data relatively well. However, the competition model resulted in a qualitatively poor 1t to the [001] construct as shown by the lack of saturation in the most anterior region of the embryo (Fig. S5C, ii). The direct repression and quenching models showed equally good 1ts to the data stemming from this construct.

## S5.3 Predicting two-Runt binding sites data for each mode of repression

We further tested these different models of repression by using the parameters inferred from the one-Runt binding site constructs to predict the rate of initiation for the two-Runt binding sites constructs. As reasoned in the main text, we began by assuming that the two Runt molecules act independently of each other such that there are no interactions between Runt molecules. Figure S6 shows this parameter-free prediction for our two-Runt binding sites constructs for all three modes of repression. As shown in the figure, none of the models can explain the data, suggesting the need to invoke additional interactions between the molecular players of our model.

Next, we considered whether Runt-Runt pairwise or higher-order cooperativities had to be invoked in order to explain the two-Runt binding sites data for both the competition and quenching mechanisms. For the competition model, we considered Runt-Runt cooperativity, *ω_rr_*, and Runt-Runt-Bicoid higher-order cooperativity, *ω_brr_* in addition to the Runt-Bicoid interaction term *ω_br_*. In the quenching scenario, we accounted for Runt-Runt cooperativity, *ω_rr_*, and Runt-Runt-Bicoid-RNAP higher-order cooperativity, *ω_brrp_*. For both the competition (Fig. S7) and quenching (Fig. S8) mechanisms, we observed a qualitatively similar trend to that observed for direct repression (Fig. 6). Specifically, as shown in Figures S7C and S8C, considering pairwise cooperativity did not signi1cantly improve the MCMC 1ts to the data for either model considered. Further, considering only the higher-order cooperativity also did not improve the 1ts for both competition and quenching mechanisms as shown in Figure S7D and Figure S8D. Invoking both Runt-Runt cooperativity and higher-order coop-erativity improved the 1ts qualitatively for both competition and quenching mechanisms as shown in Figure S7E and Figure S8E.

While the quenching model showed almost equally good MCMC 1ts to the data as the direct repression model, the competition model showed qualitatively poor 1ts in any combination of cooperativities. In particular, there was a significant mismatch in the most anterior region of the embryo, where Bicoid is thought to saturate *hunchback* expression. While we do not view these 1ts as conclusive evidence to support one mechanism over the other, an exercise that would require a new round of experimentation, we conclude that higher-order cooperativity is required to explain the data from the two-Runt binding sites constructs regardless of the choice of mechanism of Runt.

## S6 Design of synthetic enhancer constructs based on the *hunchback* P2 enhancer

The Runt binding sites were introduced into the *hunchback* P2 minimal enhancers at the positions determined by Chen et al. [2012]. To make this possible, the authors chose positions containing presumed neutral DNA sequences, meaning that these DNA locations did not contain obvious motifs for Bicoid or Zelda, the major input transcription factors that regulate this enhancer. Then, these DNA sequences were mutated to turn them into Runt binding sites.

To ensure that this process did not perturb the binding sites for Bicoid and Zelda we resorted to the Advanced PATSER entry form [Hertz et al., 1990, Hertz and Stormo, 1999] which identi1es the location of transcription factor binding sites from a sequence of DNA based on position weight matrices. We used position weight matrices for Bicoid and Zelda from Park et al. [2019]. PATSER was run with the settings described in Eck et al. [2020] for both the *hunchback* P2 enhancer and the *hunchback* P2 enchancer with three Runt binding sites (from Chen et al. [2012]) for Bicoid and Zelda, respectively. The result of this analysis for these two constructs is shown for each transcription factor in Figure S9A. Here, we took a the PATSER score cutoff—for considering a given sequence to be a binding site—of 3 as in Eck et al. [2020]. We observed that the recognized binding motifs for both Bicoid and Zelda were identical between the two constructs, meaning that we did not add additional Bicoid or Zelda binding sites by introducing the Runt motifs. The resulting synthetic enhancer with three Runt binding sites with mapped binding sites for Bicoid, Zelda (Fig. S9A), and Runt [Chen et al., 2012] is shown in Figure S9B as a reference. The position of the Runt binding sites are noted from their distance from the promoter (which is marked as 0).

## S7 Quantifying the nuclear concentration of LlamaTag-Runt

The major caveat in the eGFP:LlamaTag-Runt fluorescence measurements is that the raw nuclear fluorescence that we measured consists of two populations: eGFP *bound* to the LlamaTag-Runt, and *free*, *unbound* eGFP. Thus, in order to measure nuclear Runt concentration, we need to factor out the contribution from free eGFP to the overall fluorescence.

We followed the procedure described in Bothma et al. [2018] which consists of using cytoplasmic fluorescence to calculate the free nuclear eGFP under two assumptions. First, we posit that most of the transcription factors reside in the nucleus such that the cytoplasmic fluorescence mostly reports on free cytoplasmic eGFP. Second, we assume that the nucleus-to-cytoplasm ratio of free eGFP is kept constant at a measured chemical equilibrium of *K_G_* = *GF P_-C_* /*GF P_N_* = 0.8, where *GF P_-C_* and *GF P_N_* are the eGFP fluorescence in nuclei and cytoplasm in the absence of LlamaTag [Bothma et al., 2018].

As shown in Bothma et al. [2018], the nuclear concentration of the GFP-tagged transcription factor, *GF P* − *T F_N_*, is given by

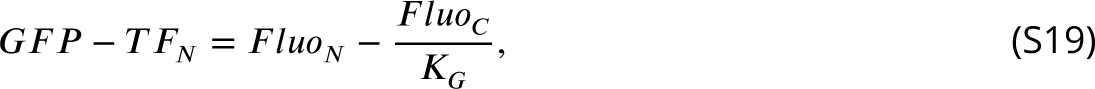

where *F luo_N_* and *F luo_-C_* are the eGFP fluorescence in nuclei and cytoplasm, respectively, that we measured in the embryos with both eGFP and LlamaTagged Runt. The resulting nuclear concentration of LlamaTag-Runt is shown in Figure 3B.

## S8 Quantitative interpretation of MS2 signals

The MS2 signal reports on three features of transcriptional dynamics: 1) the initial RNAP loading rate, 2) the duration of transcription, and 3) the fraction of loci that engage in transcription at any time point in the nuclear cycle. In this section, we will explain in further detail how we extract these features from the MS2 signal over nuclear cycle 14.

### S8.1 Extracting the initial RNAP loading rate

The initial rate of RNAP loading corresponds the average transcription rate observed after transcriptional onset and until the MS2 signal reaches its peak value during nuclear cycle 14. In order to measure this rate, we followed the protocol described in Garcia et al. [2013]. Briefly, as shown in Figure S10A, we fitted a line to the MS2 time trace (averaged over nuclei within a spatial window of 2.5% of the embryo length) within the time window of 5 to 10 minutes after the 13th anaphase. The slope of this line reported on the initial rate of RNAP loading (Fig. 3G). The spatial profiles of this initial rate of RNAP loading across all our synthetic enhancer constructs and genotypes are shown in Figure S10B.

### S8.2 Extracting the duration of transcription

In the main text, we focused on the theoretical prediction of the initial rate of transcription. However, the length of the time window over which transcription occurs [Lammers et al., 2020] is another regulatory knob that, in principle, Runt could modulate to dictate gene expression patterns. We sought to determine the duration of time over which transcription occurs to assess whether Runt affects not only the initial rate of transcription, but also the time window over which transcription could initiate. To quantify the effective duration of transcription initiation, we resorted to the analysis methodology developed in Garcia et al. [2013]. Briefly, we parametrized the MS2 signal decay regime—after transcription reaches its peak and becomes slower than the unloading rate [Garcia et al., 2013]—as an exponential decay (Fig. S11A). Thus, we can describe the MS2 spot fluorescence trace in the decay regime as

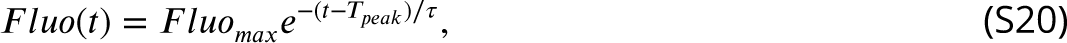

where *T_peak_* represents the time point where the MS2 spot fluorescence reaches its peaks, and *τ* is the decay time.

Given the sometimes noisy MS2 traces (data not shown), we fitted an exponential curve to the more robust integral of the MS2 spot fluorescence over time from *T_peak_* to the end of nuclear cycle 14 as shown in Figure S11B. This quantity is proportional to the amount of mRNA produced between the integration bounds [Garcia et al., 2013]. The resulting accumulated mRNA time trace is then fitted to the integrated form of Equation S20, which is given by

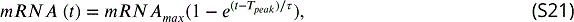

where *mRNA_max_* is the accumulated mRNA at the end of nuclear cycle 14.

The resulting profiles of the duration of transcription along the embryo for our all synthetic enhancer constructs are illustrated in Figure S11C in the presence and absence of Runt protein. As shown in the figure, this duration time is not significantly modulated by Runt repressor.

### S8.3 Calculation of the fraction of competent nuclei

Another quantity that could be modulated by Runt repressor is the fraction of loci that ever engage in transcription during a given nuclear cycle, which we termed as the “fraction of competent loci”. As demonstrated by Garcia et al. [2013], Dufourt et al. [2018], Lammers et al. [2020] and Eck et al. [2020], this fraction of transcriptionally competent loci is modulated along the anterior-posterior axis, presumably due to the action of transcription factor gradients.

To show a concrete example of how this quantity is calculated, we take data from one construct ([000]) showing the MS2 spot fluorescence time traces from individual loci of transcription as shown in Figure S12A. Here, columns represent time points during nuclear cycle 14, and rows represent individual transcriptional loci. As shown in the figure, roughly 80% of the loci, labeled as “competent loci”, show active transcription during nuclear cycle 14. However, the remaining 20% of the loci never engage in transcription, which we termed as “incompetent loci”. Because these two populations exhibit wildly different behaviors, we de1ne the fraction of competent loci as

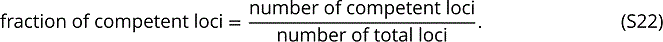

Thus, in this example in Figure S12A, the fraction of competent loci is approximately 0.8.

Figure S12B shows the measured fraction of active loci for all synthetic enhancer constructs in the presence and absence of Runt repressor. As seen in the figure, although this quantity can be modulated by the presence of Runt repressor, this is not always the case (e.g., [010] and [11]). Moreover, we could not 1nd a trend for how the fraction of competent loci is modulated by different combinations of Runt binding sites. For example, the [100] construct alone did show a change in the fraction of active loci in the presence of Runt, whereas the [010] construct did not. When these two binding sites were combined as the [110], there was no significant modulation of the fraction of competent loci when adding Runt repressor. In another example, the [001] construct showed a mild modulation of the fraction of competent loci. However, when this Runt binding site was combined with the [010], which did not show any modulation, the [011] construct showed a much bigger modulation of the fraction of competent loci than the [001]. Thus, the [010] Runt binding site could drive more or less modulation of the fraction of competent loci when combined with different Runt binding sites in a context-dependent manner. As a result of our failure to uncover an apparent trend in terms of which regulatory architectures lead to a stronger modulation of the fraction of active loci, we did not attempt to theoretically explain the regulation of this fraction of active loci in this study.

### S9 Supplementary figures

**Figure S1.**
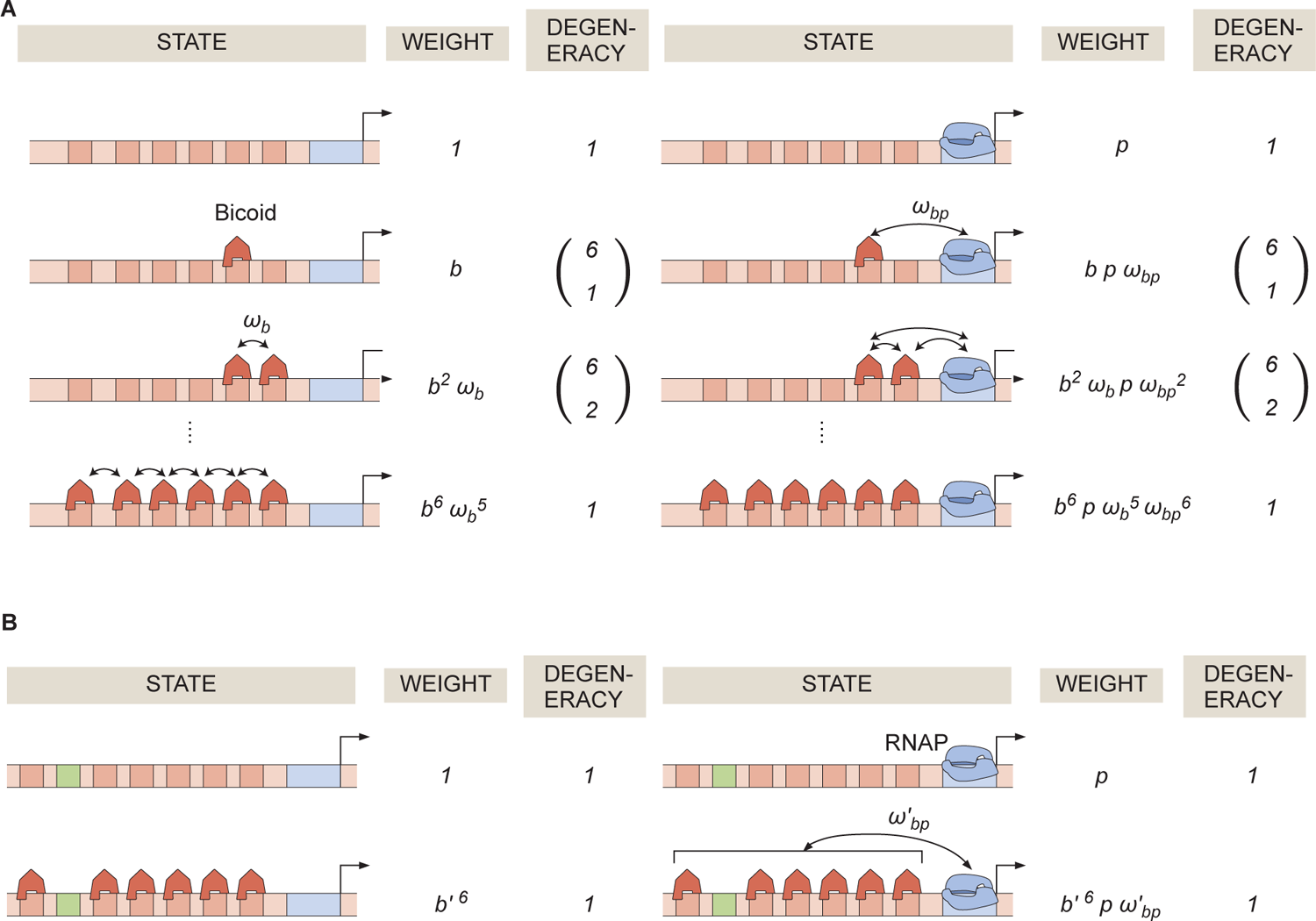
General thermodynamic model for a *hunchback* P2 enhancer with six Bicoid binding sites. **(A)** States, weights, and degeneracy considered for our thermodynamic model. **(B)** Simpler form of the thermodynamic model in the limit of *ω_b_ »* 1.

**Figure S2.**
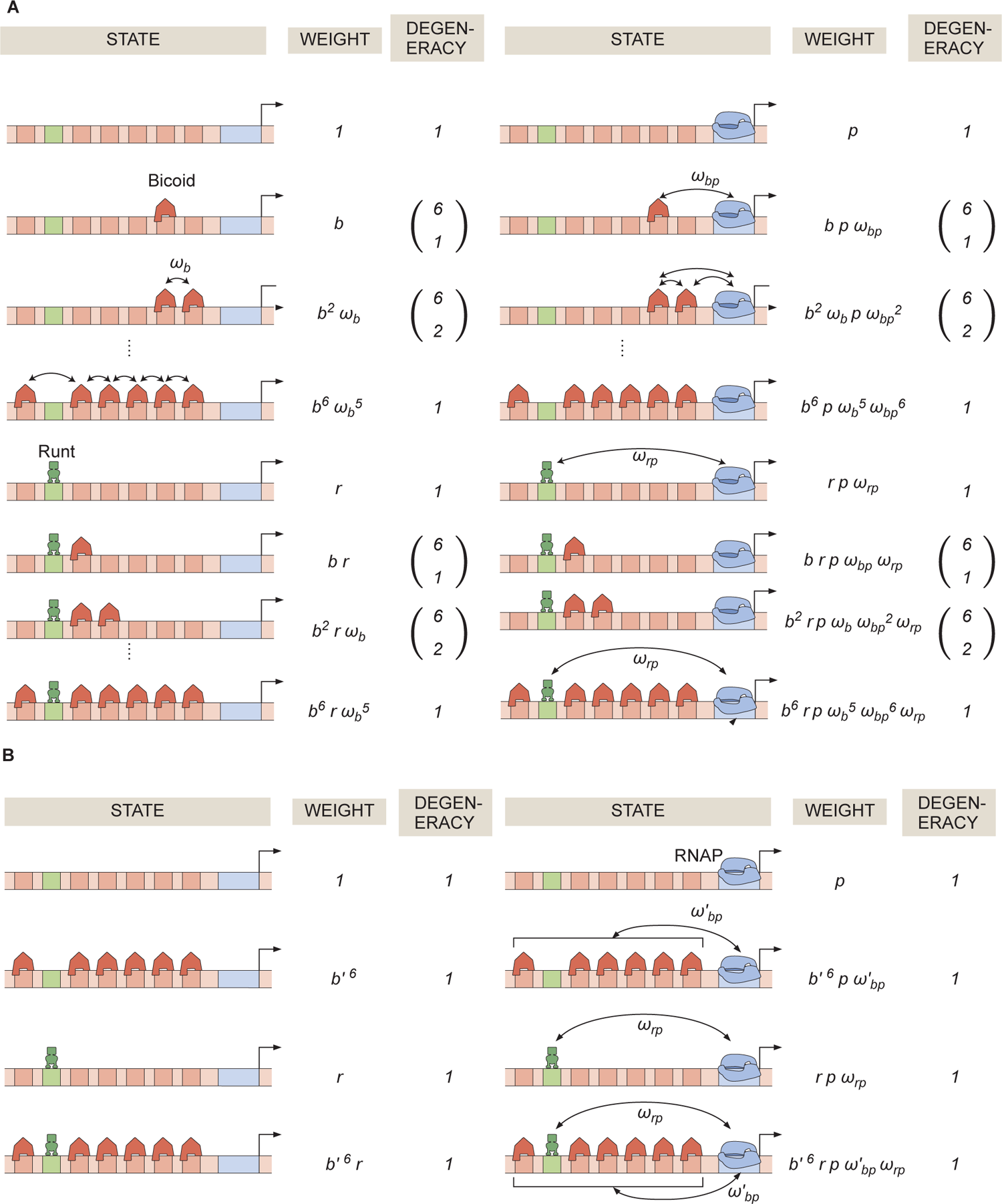
General thermodynamic model for an enhancer with six-Bicoid binding sites and one Runt binding site. **(A)** Statistical weights and degeneracy of each state the system can be found in. **(B)** Simpler form of the model from (A) in the limit of strong Bicoid-Bicoid cooperativity.

**Figure S3.**
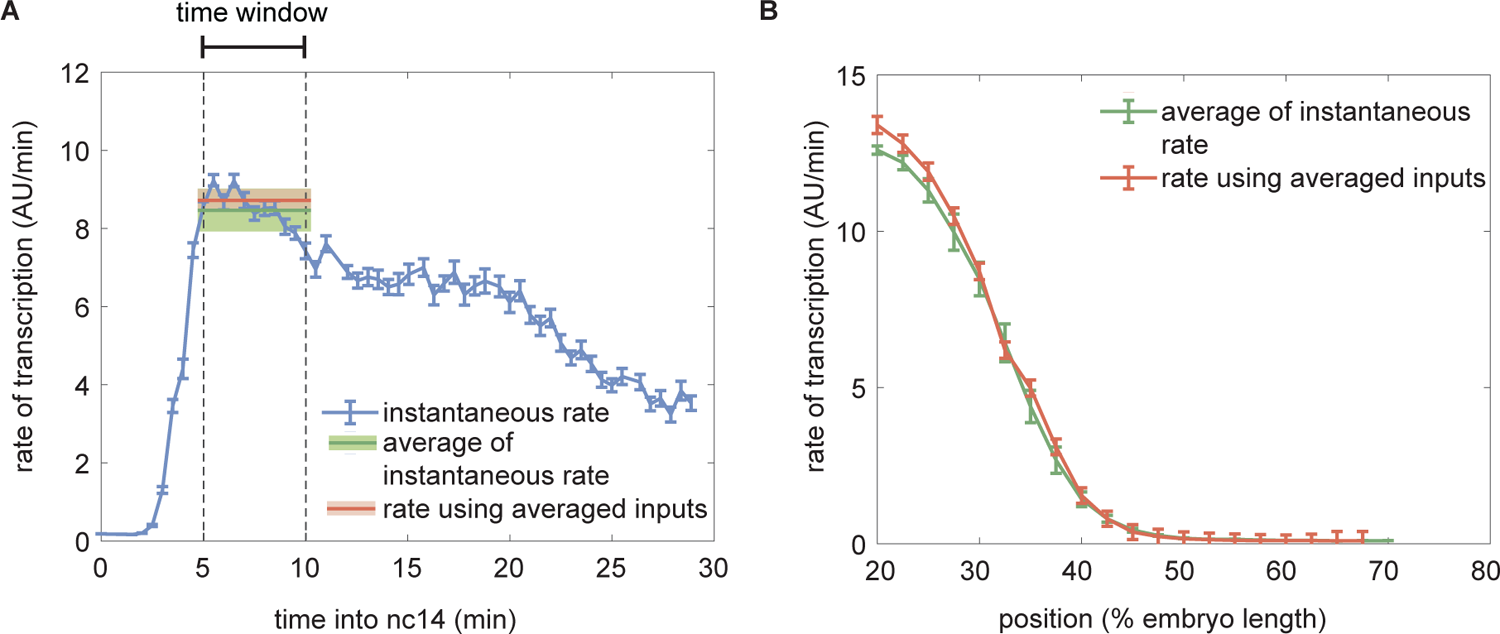
Comparison of the predicted rate of transcription using dynamic and time-averaged transcription factor concentration pro1les as inputs. **(A)** Instantaneous predicted rate of transcription calculated using dynamic transcription factor concentration pro1les at each time point (blue) and resulting averaged rate of transcription averaged over the time window of 5-10 minutes from the 13th anaphase (green) compared to the predicted rate of transcription obtained using the static transcription factor concentrations of Bicoid and Runt shown in Figure 3E (red). (Illustrative predictions calculated at 30% of the embryo length using *K_b_* = 30(*AU*), *K_r_* = 100(*AU*), *ω_bp_* = 100, *ω_rp_* = 0.1, *p* = 0.001, *R* = 300(*AU* /*min*).) **(B)** Spatial pro1le of the predicted rate of transcription calculated by averaging the instantaneous transcription rate (green) or by using the averaged input transcription factor concentrations as inputs (red). (A, B, error bars and shaded areas represent the standard error of mean over embryos 42 embryos generated from making pairs of independently measured six eGFP-Bicoid embryo and seven GreenLlamaTag-Runt embryo.)

**Figure S4.**
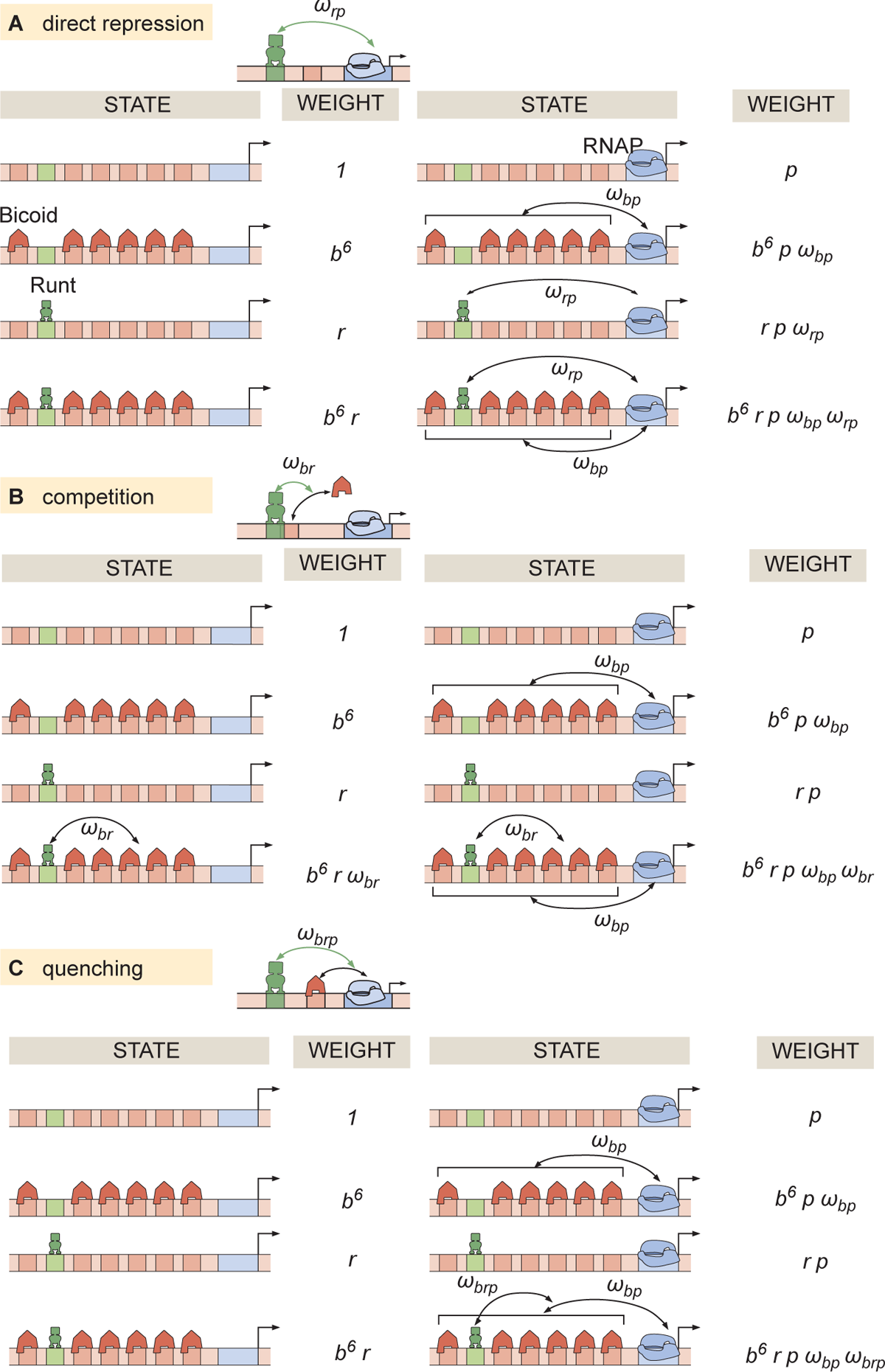
Thermodynamic models for different modes of repression. States and statistical weights corresponding to the *hunchback* P2 enhancer with one Runt binding site for the **(A)** direct repression, **(B)** competition, and **(C)** quenching mechanisms.

**Figure S5.**
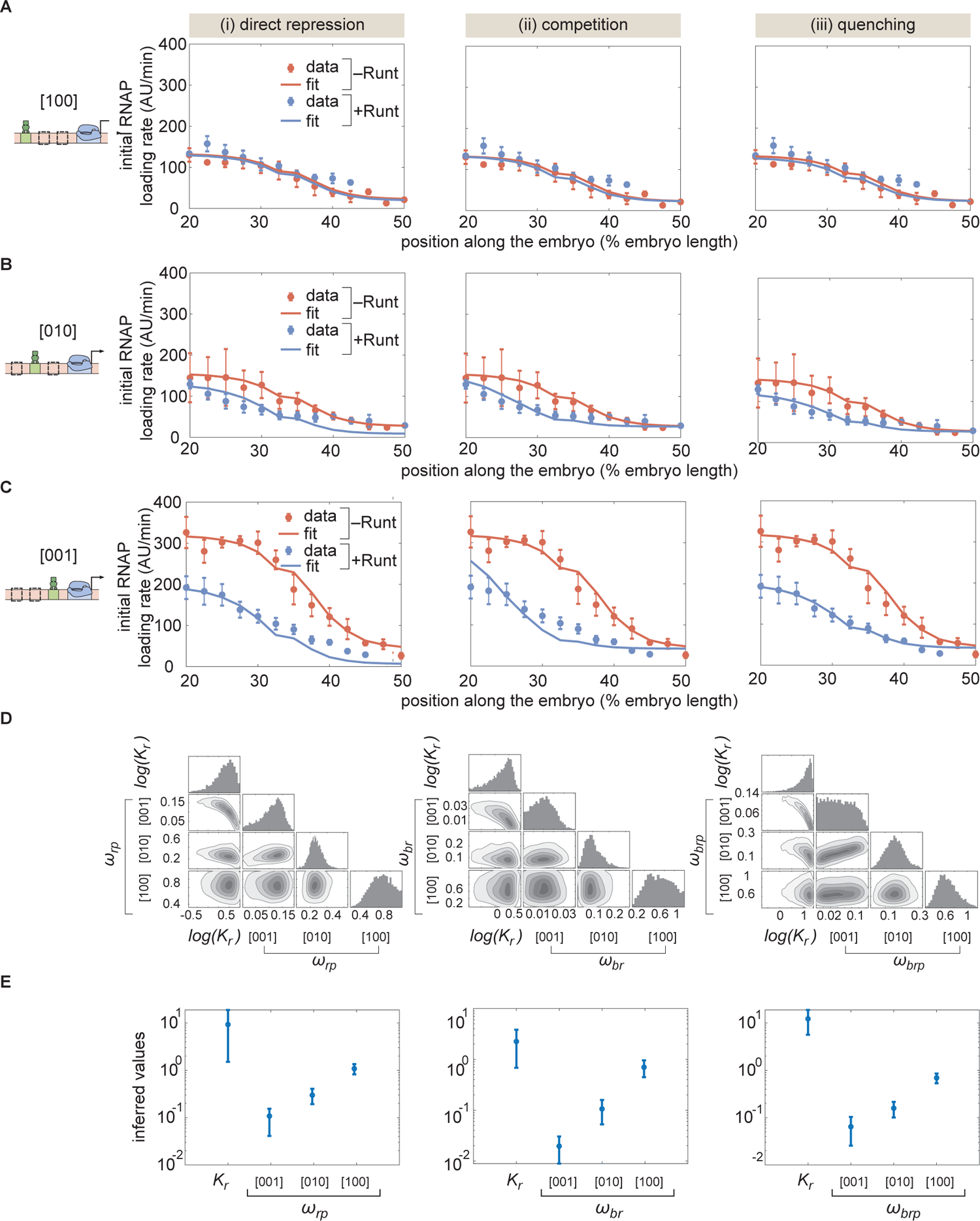
MCMC 1tting to the *hunchback* P2 with one Runt binding site constructs using different models of repression. **(A,B,C)** MCMC 1ts for three modes of repression, (i) direct repression, (ii) competition, and (iii) quenching, for our three one-Runt site constructs, **(A)** [100], **(B)** [101], and **(C)** [001]. **(D)** Corner plots resulteing from MCMC inference on the three one-Runt site constructs for each model. **(E)** Inferred parameters from MCMC 1tting. (A,B, and C, error bars represent standard error of the mean over 3 embryos; E, error bars represent standard deviation of the posterior chain.)

**Figure S6.**
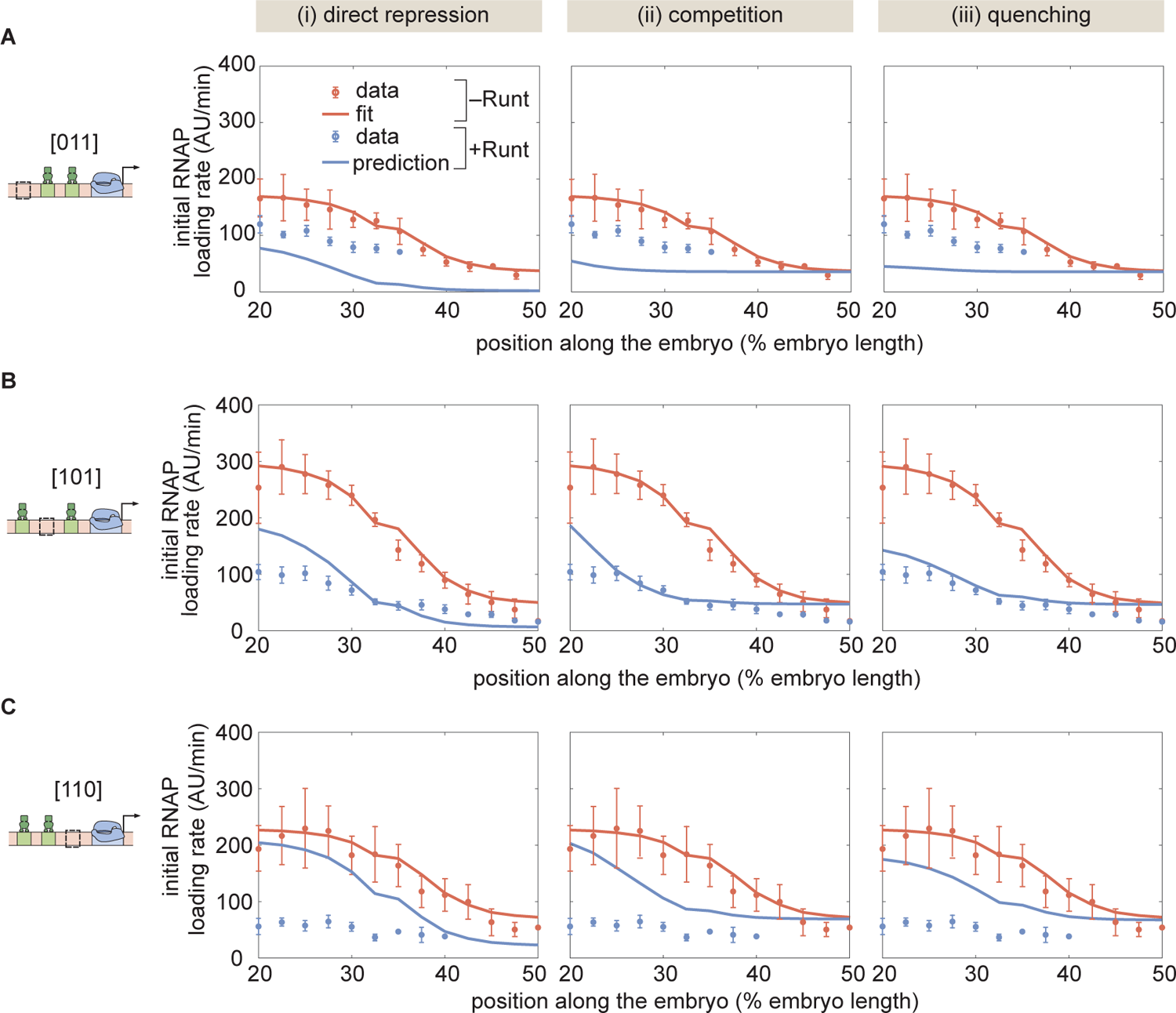
Prediction for two-Runt binding sites constructs based on the inferred parameters from the one-Runt binding site cases for different modes of repression for the **(A)** [011], **(B)** [101], and **(C)** [110] constructs. The model assumes no interactions between Runt molecules. (A,B, and C, error bars represent standard error of the mean over 3 embryos.)

**Figure S7.**
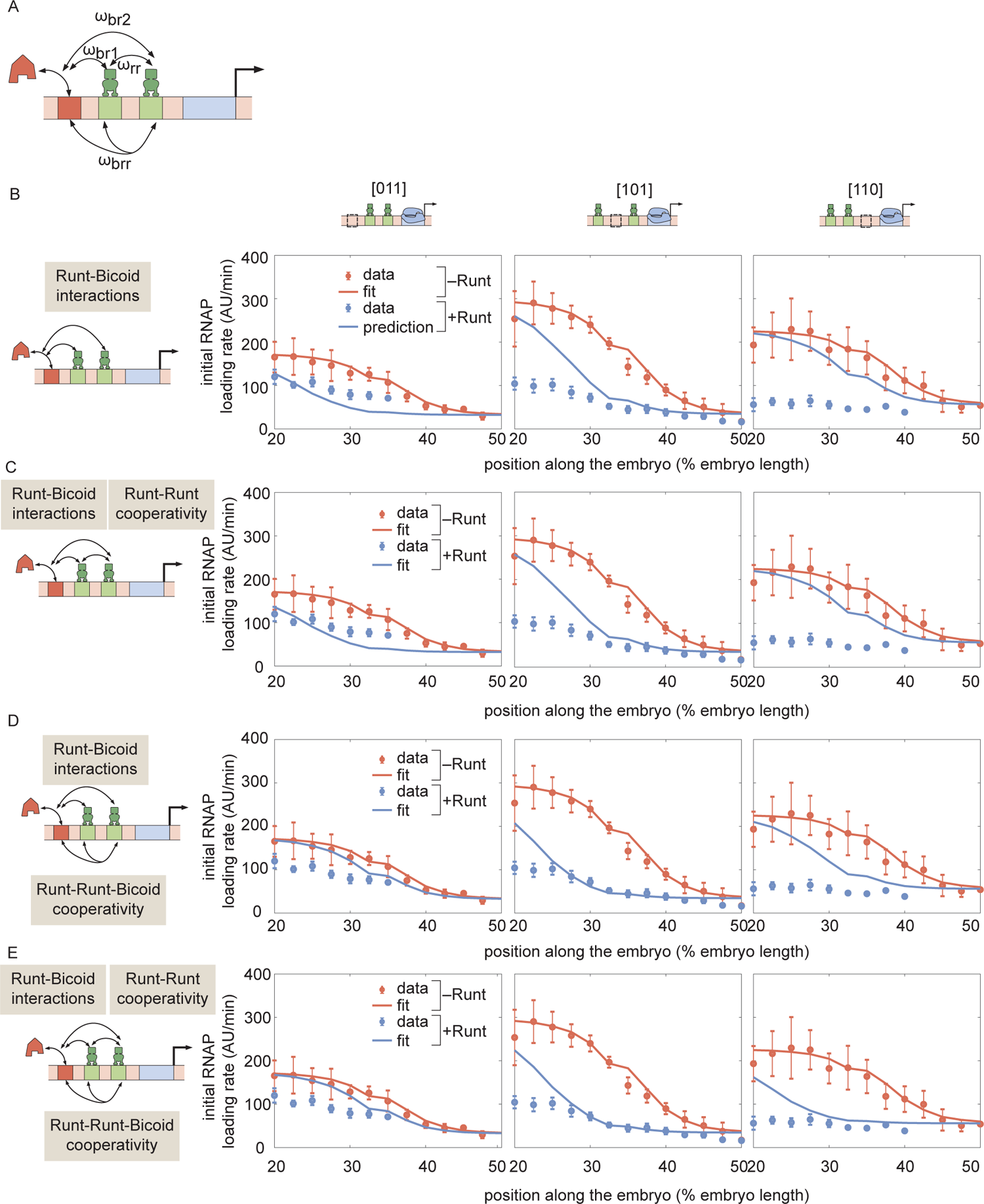
Prediction for *hunchback* P2 transcription initiation rate with two-Runt binding sites under the competition scenario for different combinations of cooperativities. **(A)** Schematic of cooperativity terms considered: Runt-Runt cooperativity given by *ω_rr_* and Runt-Runt-Bicoid complex higher-order cooperativity captured by *ω_brr_*, in addition to the competition terms *ω_br_*_1_ and *ω_br_*_2_. **(B)** Zero-parameter prediction using the inferred parameters from zero- and one-Runt binding site constructs. **(C,D,E)** Best MCMC 1ts for our three two-Runt binding sites constructs considering **(C)** Runt-Runt cooperativity, **(D)** Runt-Runt-Bicoid complex higher-order cooperativity, and **(E)** both Runt-Runt cooperativity and Runt-Runt-Bicoid complex higher-order cooperativity. (B,C,D, and E, error bars represent standard error of the mean over 3 embryos.)

**Figure S8.**
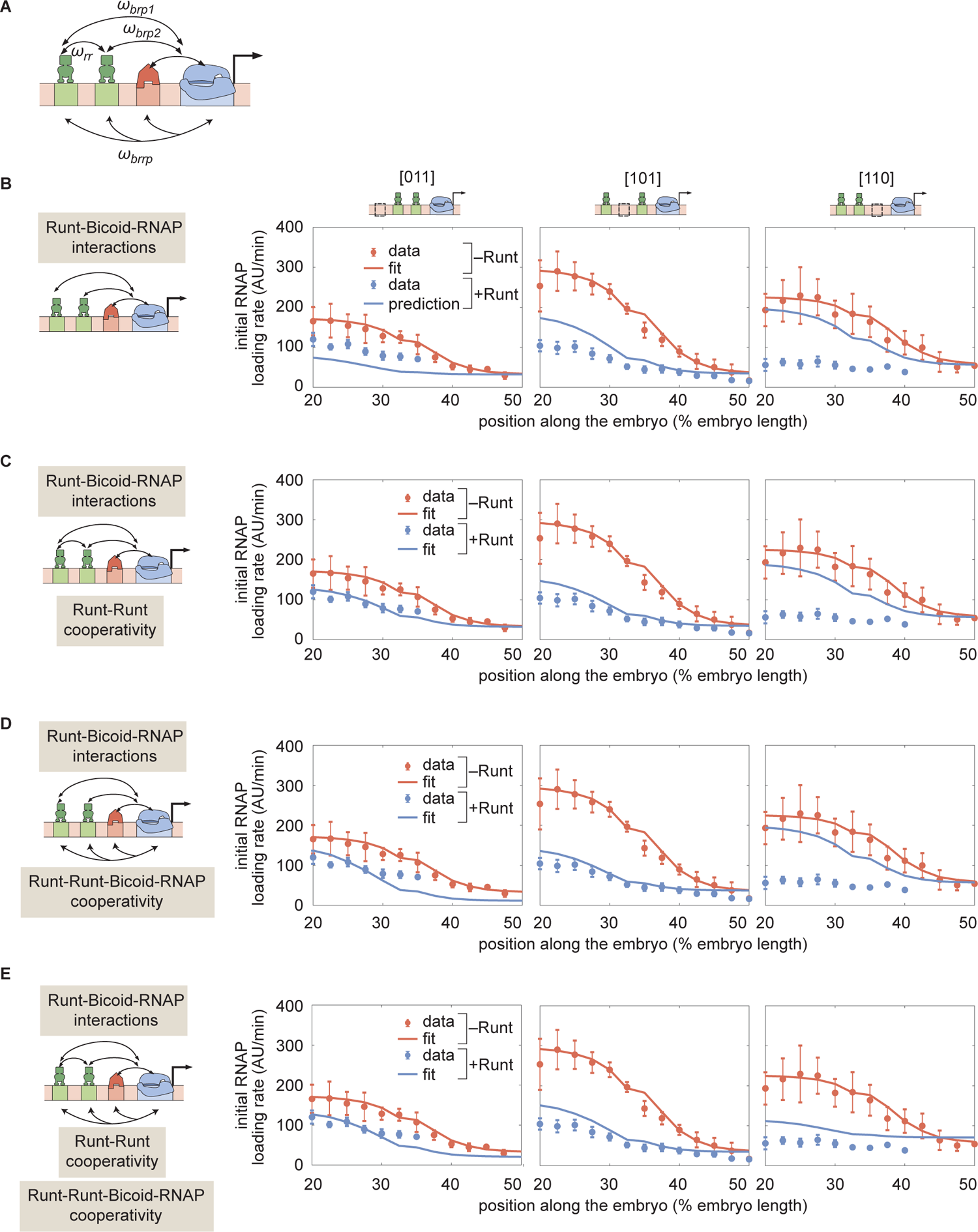
Prediction for *hunchback* P2 transcription initiation rate with two-Runt binding sites under the quenching mechanism for different combinations of cooperativities. **(A)** Schematics of additional cooperativities considered: Runt-Runt cooperativity *ω_rr_* and Runt-Runt-Bicoid-RNAP complex higher-order cooperativity *ω_brrp_*. **(B)** Zero-parameter prediction using the inferred parameters from one-Runt binding site constructs. **(C,D,E)** Best MCMC 1ts for our three two-Runt binding sites constructs considering **(C)** Runt-Runt cooperativity, **(D)** Runt-Runt-Bicoid-RNAP higher-order cooperativity, and **(E)** both Runt-Runt cooperativity and Runt-Runt-Bicoid-RNAP higher-order cooperativity. (B,C,D, and E, error bars represent standard error of the mean over 3 embryos.)

**Figure S9.**
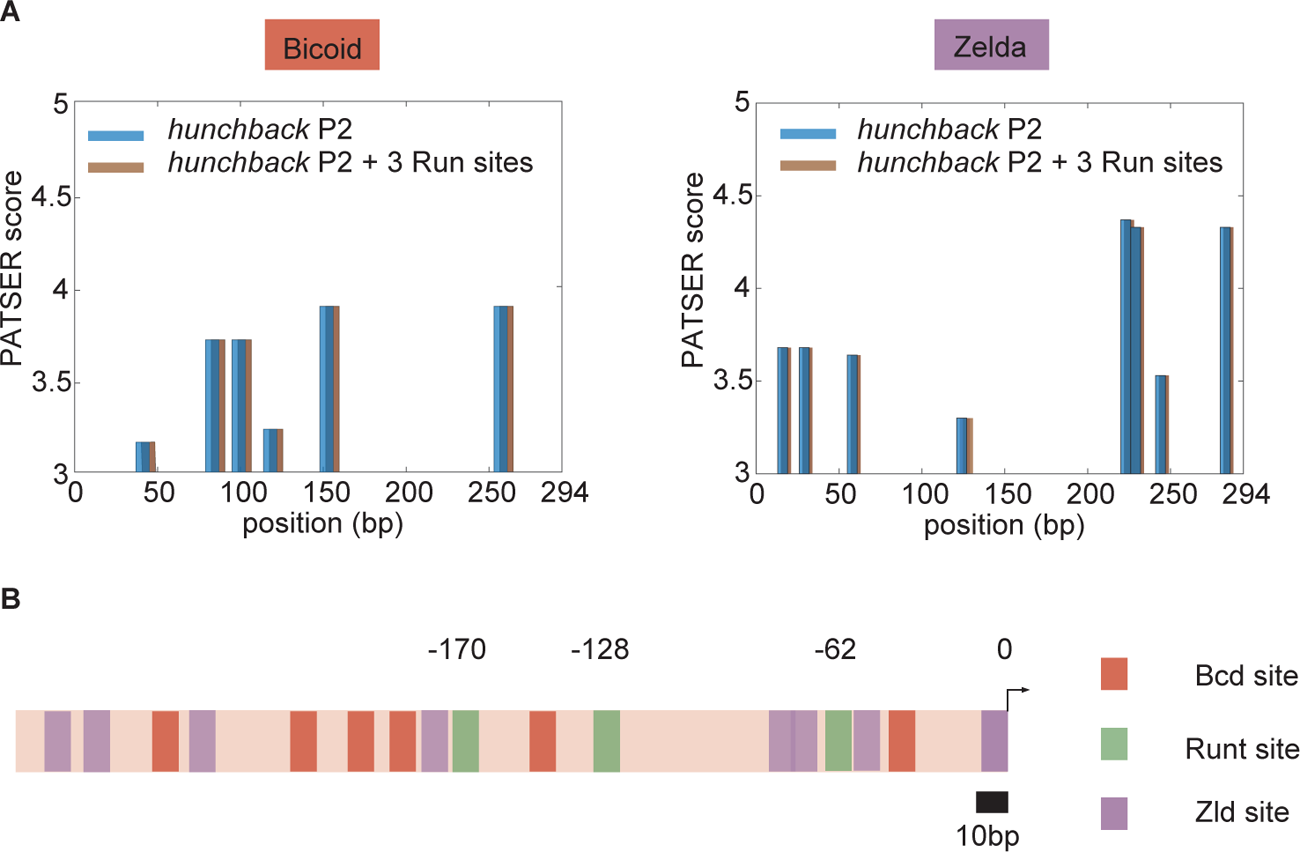
Bioinformatically predicted architecture of major transcription factor binding sites in the *hunchback* P2 minimal enhancer with three Runt binding sites. **(A)** PATSER scores for Bicoid and Zelda for *hunchback* P2 (blue) and *hunchback* P2 with three Runt sites (brown). The binding motifs with PATSER scores higher than three are shown. We concluded that neither Bicoid nor Zelda binding sites were created or removed by the introduction of these three Runt binding motifs. **(B)** A schematic diagram of *hunchback* P2 minimal enhancer with three Runt binding sites with mapped binding sites for Bicoid and Zelda from (A) and Runt binding sites from Chen et al. [2012]. The position of Runt binding sites are noted with their distance from the promoter (marked as 0).

**Figure S10.**
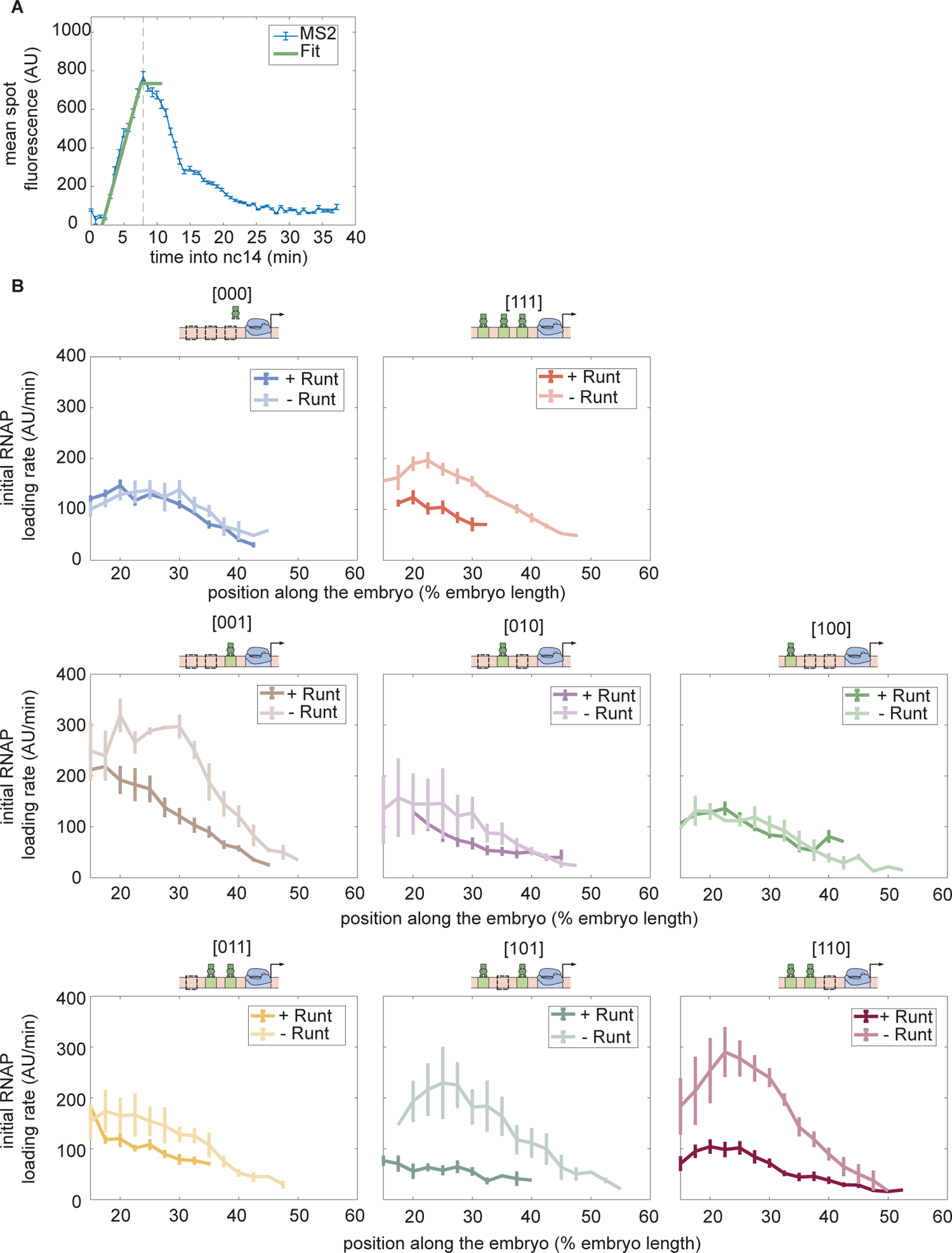
Initial rate of RNAP loading in nuclear cycle 14 across the anterior-posterior axis for different constructs, with or without Runt protein. **(A)** Schematic showing how the initial rate of RNAP loading is measured by extracting the slope resulting from a linear 1t to the MS2 time traces at the beginning of nuclear cycle 14. **(B)** Initial rate of RNAP loading along the embryo length for each construct in the presence and absence of Runt for each of our synthetic enhancer construct. (B, Error bars represent standard error of the mean over 3 embryos.)

**Figure S11.**
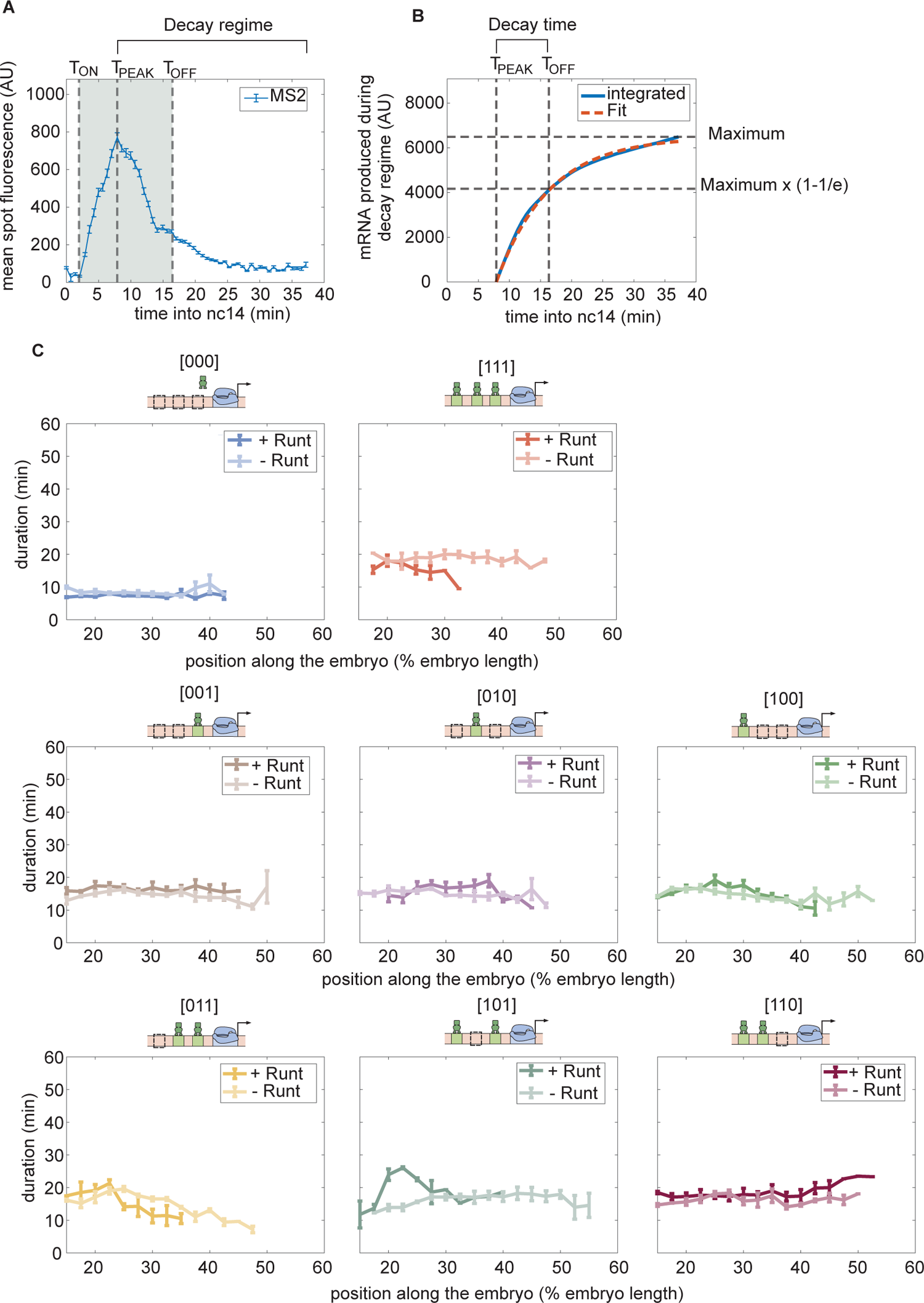
Duration of transcription over nc14. **(A)** An example MS2 time trace in nuclear cycle 14. The decay regime is de1ned from the peak of the signal to the end of the measurement. *T_ON_* is de1ned by the x-intercept of the slope of the 1tted line. *T_off_* is determined by the decay time in the exponential function. The gray shaded region from *T_ON_* to *T_OF F_* is de1ned as the transcriptional time window. **(B)** The decay time can be extracted from the accummulated mRNA signal obtained by integrating the MS2 fluorescence. Here, decay time is de1ned as the time it takes to reach (1-1/e) of that maximum accumulated mRNA. **(C)** Transcriptional time window along the anterior-posterior axis for each construct with and without Runt protein. (A, error bars represent standard error of the mean over the spatial averaging corresponding to roughly ten nuclei; C, error bars represent standard error of the mean over 3 embryos.)

**Figure S12.**
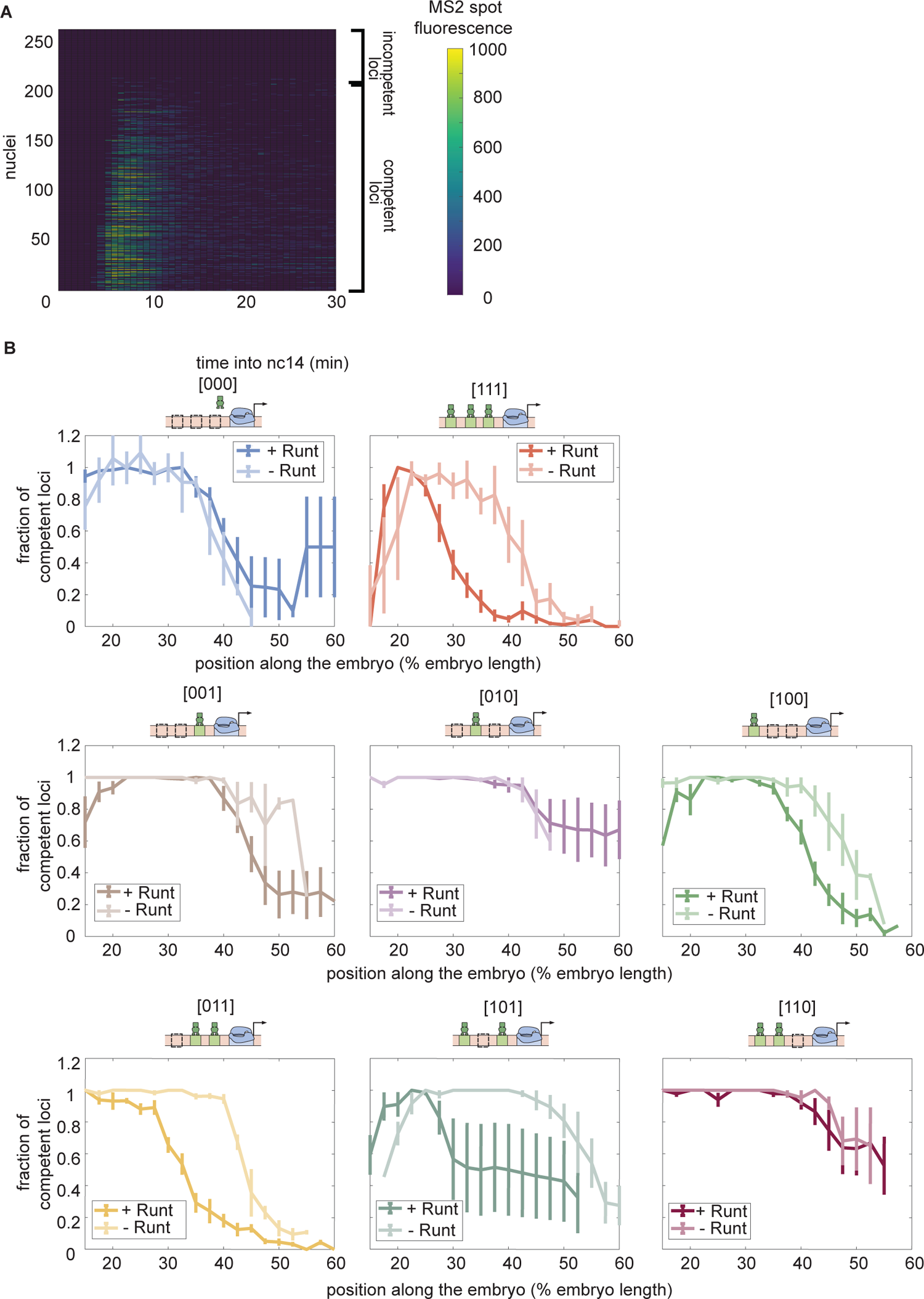
Fraction of competent loci in nuclear cycle 14 along the anterior-posterior axis for each synthetic enhancer construct in the presence and absence of Runt protein. **(A)** Heatmap showing the transcriptional signal from the *hunchback* P2 enhancer for individual nuclei (rows) demonstrating that there are two populations of loci: transcriptionally active and inactive loci. **(B)** Fraction of transcriptionally active loci along the embryo for each construct for wild-type and *runt* null backgrounds. (B, error bars represent standard error of the mean over 3 embryos.)

**Figure S13.**
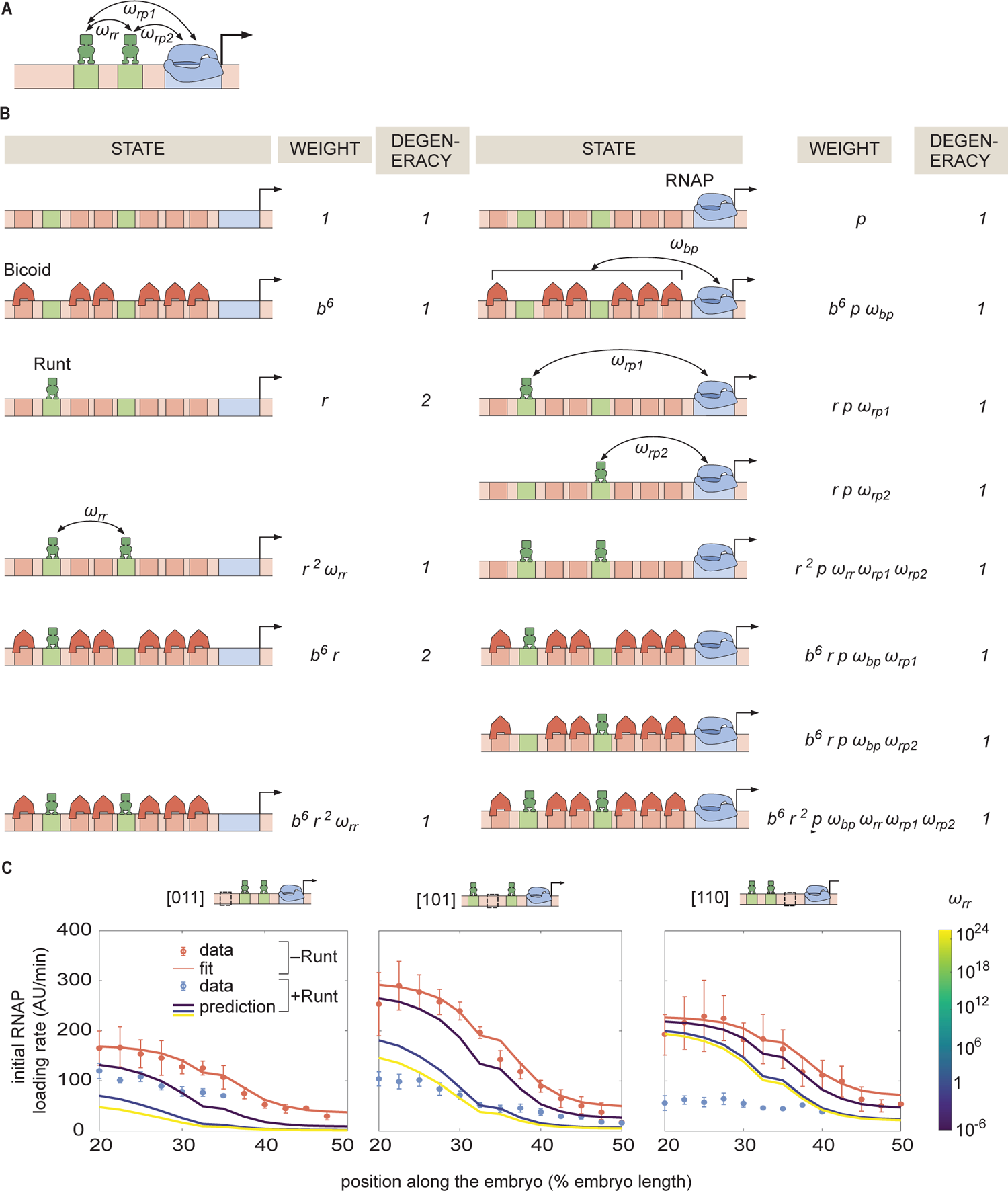
Invoking Runt-Runt cooperativity in the thermodynamic model is not sufficient to explain the experimental data from *hunchback* P2 with two Runt binding sites. **(A)** Model schematics where we add a new *ω_rr_* parameter representing Runt-Runt cooperativity. **(B)** Corresponding states and weights for *hunchback* P2 with two Runt binding sites in the presence of Runt-Runt cooperativity. **(C)** Prediction of the initial rate of RNAP loading pro1les over a range of Runt-Runt cooperativity strength, *ω_rr_* = [10^−6^, 10^24^], for all constructs of *hunchback* P2 with 2 Runt binding sites with different con1gurations. (Left) [011], (Center) [101], (Right) [110]. (C, error bars represent standard error of the mean over 3 embryos)

**Figure S14.**
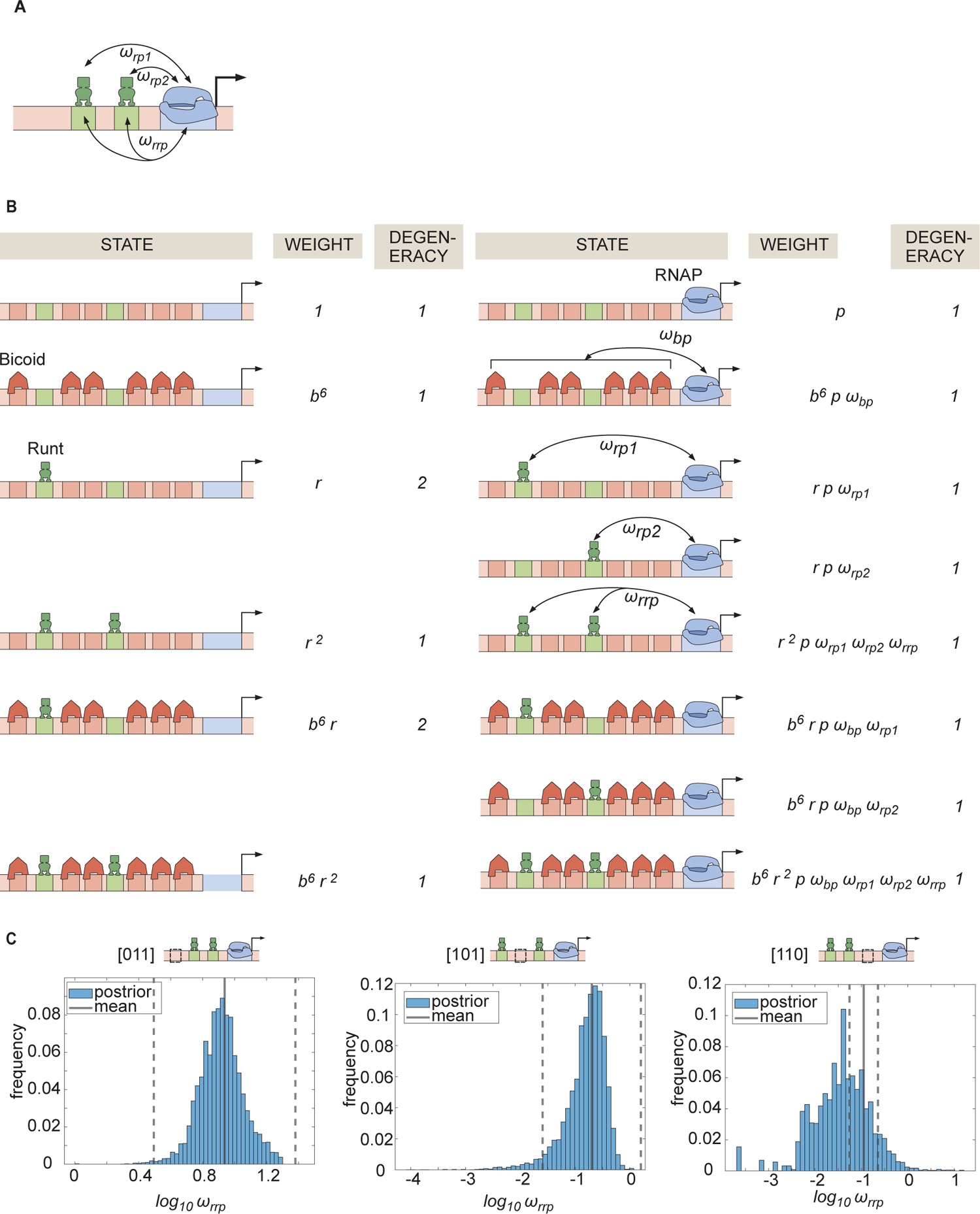
Invoking Runt-Runt-RNAP higher-order cooperativity is not sufficient to explain the two-Runt sites data. **(A)** Schematics of a model where we add Runt-Runt-RNAP higher-order cooperativity represented by *ω_rrp_*. **(B)** Thermodynamic model states and weights for *hunchback* P2 with two Runt binding sites in the presence of Runt-Runt-RNAP higher-order cooperativity. **(C)** Histograms showing the posterior distribution of the inferred *ω_rrp_* parameter from the best MCMC 1t shown in Figure 6D. The black line represents the mean and the dotted lines represent standard deviation from the mean.

**Figure S15.**
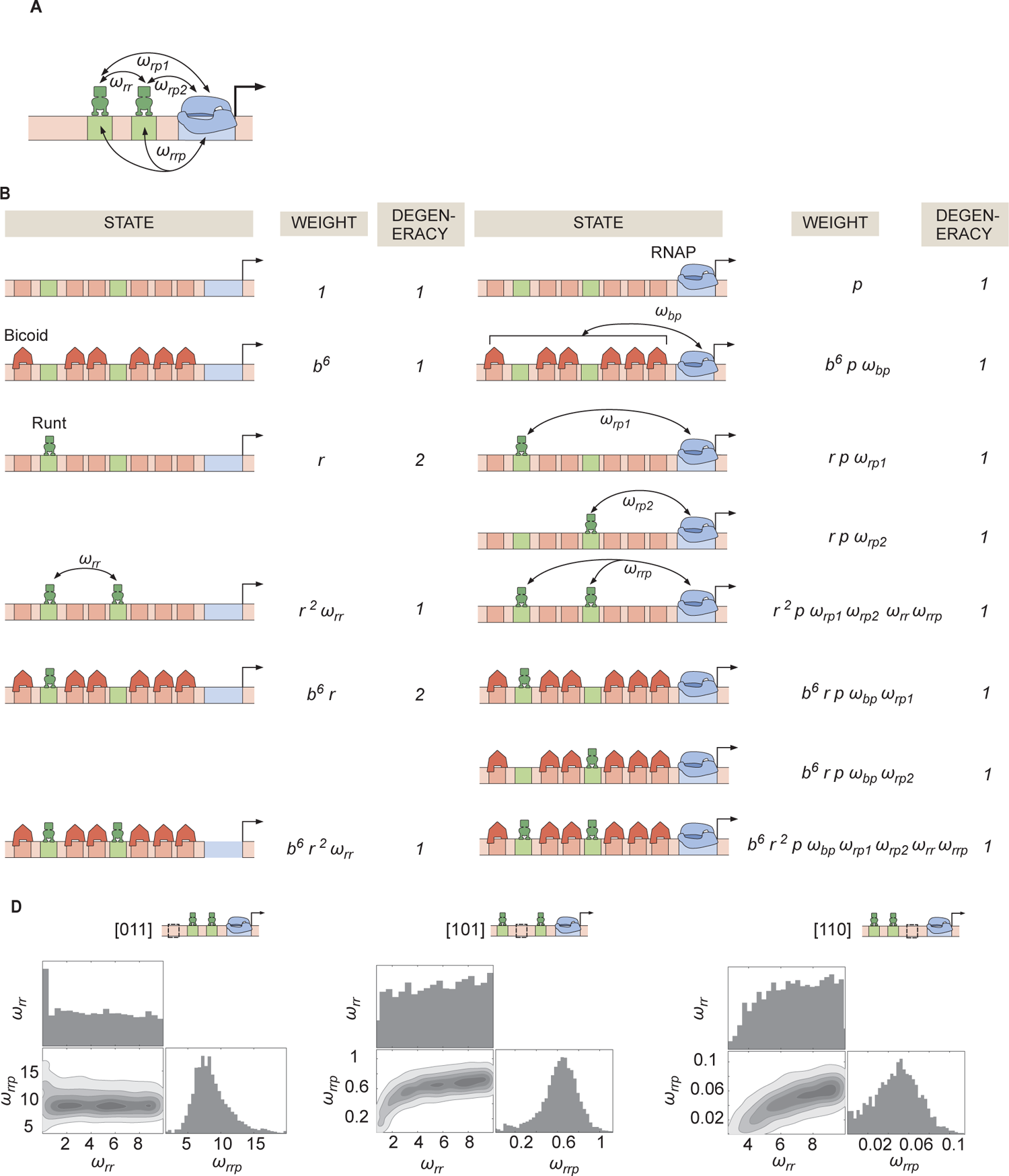
Invoking Runt-Runt cooperativity and higher-order cooperativity can explain the experimental data from *hunchback* P2 with two Runt binding sites. **(A)** Schematic showing Runt-Runt cooperativity and higher-order cooperativity. **(B)** States and weights for *hunchback* P2 with two Runt binding sites with Runt-Runt cooperativity and higher-order cooperativity. **(C)** Corner plots associated with the MCMC inference performed on two-Runt binding sites data from the best MCMC 1t shown in Figure 6E. While *ω_rr_* is not very well constrained, *ω_ho_* shows a unique optimal value.

### S10 Supplementary videos

For better quality of visualization, we recommend downloading these videos.

S1. Video S1. **eGFP-Bicoid confocal movie.** Confocal microscopy movie taken on a developing fly embryo (*eGFP-Bicoid; His2Av-mRFP; +*) during nuclear cycle 13 and 14.

S2. Video S2. **eGFP:LlamaTag-Runt confocal movie.** Confocal microscopy movie taken on a developing fly embryo (*eGFP-Bicoid; His2Av-mRFP; +*) during nuclear cycle 13 and 14.

S3. Video S3. **[001]-MS2V5:MCP-GFP (+Runt) confocal movie.** Confocal microscopy movie taken on a developing fly embryo (yw; His2Av-mRFP; MCP-eGFP) for the [001] construct with MS2 reporter during nuclear cycle 13 and 14.

